# RGS10 Attenuates Systemic Immune Dysregulation Induced by Chronic Inflammatory Stress

**DOI:** 10.1101/2024.10.24.620078

**Authors:** Janna E. Jernigan, Hannah A. Staley, Zachary Baty, MacKenzie L. Bolen, Beatriz Nuñes Gomes, Jenny Holt, Cassandra L. Cole, Noelle K. Neighbarger, Kruthika Dheeravath, Andrea R. Merchak, Kelly B. Menees, Stephen A. Coombes, Malú Gámez Tansey

**Author notes:** Corresponding Author: Dr. Malú Gámez Tansey, Department of Neuroscience, University of Florida, College of Medicine, Gainesville, FL, USA.

## Abstract

Regulator of G-protein signaling 10 (RGS10), a key homeostatic regulator of immune cells, has been implicated in multiple diseases associated with aging and chronic inflammation including Parkinson’s Disease (PD). Interestingly, subjects with idiopathic PD display reduced levels of RGS10 in subsets of peripheral immune cells. Additionally, individuals with PD have been shown to have increased activated peripheral immune cells in cerebral spinal fluid (CSF) compared to age-matched healthy controls. However, it is unknown whether CSF-resident peripheral immune cells in individuals with PD also exhibit decreased levels of RGS10. Therefore, we performed an analysis of RGS10 levels in the proteomic database of the CSF from the Michael J. Fox Foundation Parkinson’s Progression Markers Initiative (PPMI) study. We found that RGS10 levels are decreased in the CSF of individuals with PD compared to healthy controls and prodromal individuals. Moreover, we find that RGS10 levels decrease with age but not PD progression and that males have less RGS10 than females in PD. Importantly, studies have established an association between chronic systemic inflammation (CSI) and neurodegenerative diseases, such as PD, and known sources of CSI have been identified as risk factors for developing PD; however, the role of peripheral immune cell dysregulation in this process has been underexplored. As RGS10 levels are decreased in the CSF and circulating peripheral immune cells of individuals with PD, we hypothesized that RGS10 regulates peripheral immune cell responses to CSI prior to the onset of neurodegeneration. To test this, we induced CSI for 6 weeks in C57BL6/J mice and RGS10 KO mice to assess circulating and CNS-associated peripheral immune cell responses. We found that RGS10 deficiency synergizes with CSI to induce a bias for inflammatory and cytotoxic cell populations, a reduction in antigen presentation in peripheral blood immune cells, as well as in and around the brain that is most notable in males. These results highlight RGS10 as an important regulator of the systemic immune response to CSI and implicate RGS10 as a potential contributor to the development of immune dysregulation in PD.

## Background

Parkinson’s disease (PD) is the second most common neurogenerative disease, characterized by the progressive loss of dopaminergic neurons in the substantia nigra. PD impacts many systems in the body as evidenced by the development of disordered movement, autonomic dysfunction, affective disorders, and cognitive impairment [1, 2]. There is currently no cure for PD and therapeutic options are limited to symptomatic relief, highlighting the crucial need for further research into disease physiology and novel treatment development [3]. The clinical relevance of the immune system in PD has become distinctly evident over the past three decades, starting with the identification of neuroinflammation in the brains of PD patients [4–7]. Further investigation has revealed growing evidence that the central and peripheral immune systems contribute to the pathophysiology of PD [8–10]. Specifically, in PD clinical data have revealed a plethora of peripheral immune alterations, such as an increase of pro-inflammatory cytokines and pro-inflammatory immune cell subsets in circulation and cerebral spinal fluid (CSF) and a rise in central nervous system (CNS)-infiltrating peripheral immune cells [9, 11–14].

Importantly, the CSF acts as an important niche of peripheral immune cells endogenous to the CNS [15, 16]. Peripheral immune cells within the in the CSF live within the subarachnoid space in the meninges, separated from the brain parenchyma by the pia mater [17]. The meninges can support a robust inflammatory response capable of spreading to the parenchyma via either direct infiltration of immune cells or through size exclusionary permeability of cytokines [17]. Under healthy conditions, a sample of CSF contains very few peripheral immune cells, however a recent study has demonstrated an increase in activated peripheral immune cells in the CSF of individuals with PD compared to healthy controls [14]. The shift to CSI and the corresponding peripheral immune cell population dynamic changes in the CNS may represent an early feature in PD capable of inducing neuroinflammatory conditions adept at initiating neurodegeneration[18–21].

Regulator of G-protein signaling 10 (RGS10) is a key homeostatic regulator of immune cells and has been identified to negatively regulate inflammatory responses through several pathways such as suppression of nuclear factor kappa-light-chain-enhancer of activated B cells (NF-κB) activity, stromal interaction molecule 2, and glycolytic production of reactive oxygen species (ROS) in myeloid cells[22–28]. Consistently, RGS10 is most highly expressed in immune cells and lymphoid tissue, and this is especially true of myeloid cells[29, 30]. Studies have consistently shown that loss of RGS10 in myeloid cells results in hyperinflammatory conditions which can increase tissue damage[31–34]. Moreover, RGS10 is reciprocally regulated by inflammatory signaling cascades that stimulate histone deacetylation and epigenetically reduce RGS10 at the transcriptional level[35]. Therefore, we hypothesize that chronic systemic inflammation (CSI) reduces RGS10-mediated suppression of pro-inflammatory responses by the innate immune system perpetuating a feed forward loop of increasing and sustained inflammation in the periphery.

Notably, RGS10 has been implicated in multiple diseases associated with aging and chronic inflammation including PD[36]. Preclinical studies have revealed that a neuroprotective role of RGS10 in chronic inflammation and nigrostriatal neurodegeneration[22, 23, 37]. Specifically, RGS10 deficiency coupled with CSI was sufficient to induce loss of nigral dopaminergic neurons, while introduction of RGS10 into the brain of a hemi-parkinsonian rat model was sufficient to prevent the loss of nigral dopaminergic neurons and neuroinflammation[22, 37]. Moreover, there is evidence that RGS10 deficiency can shift peripheral immune cell dynamics in the CNS in experimental autoimmune encephalomyelitis (EAE) and subthreshold PD models[23, 38]. This is especially relevant, considering the documented decrease in RGS10 levels in subsets of circulating peripheral immune cells, including non-classical and intermediate monocytes as well as CD4+ T cells, in individuals with PD[39]. To understand if peripheral immune cells in the CNS also show a decrease in RGS10 and may therefore impact the induction of neuroinflammation, we investigated the protein level of RGS10 in the CSF of healthy controls, prodromal individuals with increased risk of developing PD, and individuals with diagnosed PD in the Michael J. Fox Foundation for Parkinson’s Research Parkinson’s Progression Markers Initiative (PPMI) study. Here we demonstrate a decrease in RGS10 in the CSF of individuals with PD compared to healthy controls and prodromal individuals. Moreover, we find that RGS10 levels are also significantly predicted by age and sex but not disease progression as measured by the total UPDRS score.

Importantly, CSI has been shown to increase the risk of developing neurodegenerative diseases such as PD [20]. Common sources of CSI include: a high-fat high-fructose diet, gut dysbiosis, physical inactivity, stress, sleep disturbances or deficiency, exposure to industrial toxicants, pre-existing chronic inflammatory diseases (CIDs) (such as rheumatoid arthritis (RA), inflammatory bowel disease (IBD), irritable bowel syndrome (IBS), and psoriasis), genetics, and aging[20]. Interestingly, sources of CSI overlap significantly with risk factors for developing PD, suggesting that CSI may be a common downstream process in the development and progression of PD[19, 40, 41]; however, the role of peripheral immune cell dysregulation in this process is not known. As we and others have found decreased levels of RGS10 in the CSF and circulating peripheral immune cells of individuals with PD, we hypothesize that RGS10 regulates peripheral immune cell responses to CSI prior to the onset of neurodegeneration[39]. To test this, we induced CSI through biweekly intraperitoneal (IP) injections of low dose (1X106EU/kg) lipopolysaccharide (LPS) for 6 weeks in 5-7-month-old male and female C57B6/J mice and RGS10 KO mice to investigate peripheral immune cell responses both in the circulation as well as in and around the brain. Here, we find that RGS10 deficiency synergizes with CSI, inducing a bias towards inflammatory myeloid cells and cytotoxic cell populations, as well as a reduction in innate and adaptive crosstalk through major histocompatibility complex class II (MHCII) in the peripheral blood mononuclear cells (PMBCs) as well as CNS-associated immune cells, most notably in males. These results, for the first time, highlight RGS10 as an important regulator of the systemic immune response to CSI and implicate RGS10 as a potential contributor to the development of peripheral immune dysregulation in the pathophysiology of PD.

## Methods

### Human Data

Data from Project 151 of the Parkinson’s Progression Markers Initiative (PPMI) conducted through the Michael J. Fox Foundation for Parkinson’s Research (MJFF) was used for this study. All patients that participated in the PPMI study signed an informed consent form and this study was approved by the Institutional Review Board (IRB) at the University of Florida. For this study, there are 1158 participants; 617 of which were a part of the PD cohort; 355 were part of the prodromal PD cohort, 186 were part of the control cohort. All patients in the PD cohort must have received a Parkinson’s clinical diagnosis with a positive DAT scan. Prodromal individuals are at risk for developing PD as determined by biomarkers, genetics, and clinical features. Healthy controls have no neurological disorder, no direct relative with PD, and a normal DAT scan. CSF isolation was performed using lumbar punctures as previously described [42]. All data points from participants included in the proteomics study were recorded at the beginning of the PPMI study. The proteomics study was recorded in relative fluorescence units (RFU), a measure used to indicate the levels of protein expression in a sample of CSF. This study utilized a protein quantitative trait loci (pQTL) analysis to generate the proteomics dataset. A SOMAlogic assessment, SOMAscan, was used to perform pQTL in this case. SOMAscan is an assay capable of performing large scale proteomics. For the SOMAscan, the data was subjected to 4 stages of normalization: hybridization normalization, plate scaling, median signal normalization, and calibration. Then the data was filtered to exclude non-human SOMAmers (the molecule that tags the target proteins). The final step was to log2 transform the RFU data.

### Animals

Mice were housed in the McKnight Brain Institute vivarium at the University of Florida and maintained on a 12:12 light–dark cycle with *ad libitum* access to water and standard rodent diet chow. All animal procedures were approved by the Institutional Animal Care and Use Committee at the University of Florida and followed the *Guide for the Care and Use of Laboratory Animals* from the National Institutes of Health. Generation of the RGS10 knock out (KO) line onto a C57B6/J background has been previously described [22]. Male and female C57BL6/J (B6) and RGS10 KO (KO) mice were aged to 5-7 months old and were injected with either 1X10^6^ EU/kg of lipopolysaccharide (LPS) (*Escherichia Coli* 0111:B4, Sigma Aldrich, L2630) or sterile saline intraperitoneally (IP) twice a week for 6 weeks (10 animals/group) (**Supplemental Figure 2A**). Mice were weighed and IP injections were performed on alternate sides of the mouse for each injection to reduce scar tissue formation. Animals were given access to a water bottle and wet chow 1 day prior to injections and for the duration of the paradigm. Mice that exhibited signs of dehydration were given a subcutaneous injection of sterile saline up to 1mL/day. 4 male animals in cohort 1 did not receive a single dose of LPS in week 3 due to concerns about exceeding weight loss parameters set by IACUC. At endpoint, 24h post the last injection, mice were heavily anesthetized with isoflurane and euthanized via diaphragm laceration. Blood was extracted from the right atrium of the heart and collected in an EDTA-containing vacutainer tube. Brains were removed rapidly from the skull and hemisected, with the left hemisphere immediately processed for brain immune cell isolation, while the other hemisphere was flash frozen in liquid nitrogen and stored in -80°C. In a second cohort of mice, male C57BL6/J and RGS10 KO mice were aged to 5-7 months old and underwent the same injection regimen as the previous cohort, with either 1X10 EU/kg of LPS or sterile saline IP injections twice a week for 6 weeks (5 animals/group)(**Supplemental Figure 2D**). At endpoint, 24h post the last injection, mice were heavily anesthetized with isoflurane and euthanized via diaphragm laceration. Blood and the brain were processed for PBMC and brain immune cell isolations, respectively, which were then immediately processed for RNA extraction. RNA was stored in -80° C prior to quality control and Nanostring analysis.

### PBMC isolation

Blood was processed for PBMC isolation via ACK lysis. Briefly, 5mL of ACK lysis buffer was mixed to 250µL of blood was on ice and incubated for 5 minutes and quenched with 5mL of HBSS-/-. Samples were then centrifuged at 4°C for 5 minutes at 350x*g*. Supernatant was then removed, and the remaining pellet washed in another 5mL of HBSS-/-. Samples were centrifuged again at 4°C for 5 minutes at 350xg, supernatant removed, and pellet resuspended in 200µL of 1XPBS on ice. PBMCs were processed for flow immediately and analyzed on the FACs Diva Symphony cytometer.

### Brain Immune Cell Isolation

Immune cells from the brain were isolated using Miltenyi Biotec’s Adult Brain Dissociation Kit (ABDK, #130–107-677) according to the manufacturer’s protocol as described previously [43]. Briefly, brains were harvested, washed, and cut into approximately 16 small pieces, and put into gentleMACS C-tubes (Miltenyi Biotec, #130-093-237) with dissociation enzymes prepared for initial tissue homogenization using the 37C_ABDK_01 protocol on the gentleMACS Octo-Dissociator with heaters (Miltenyi Biotec, #130-096-427) for dissociation. Brain lysates were then filtered through a 70µM filter with Dulbecco’s PBS with calcium, magnesium, glucose, and pyruvate (D-PBS) and pelleted at 300xg for 10min at 4°C and debris and red blood cells were removed from the sample. Once cells had been cleaned, immune cells were isolated using magnetic separation with anti-CD45 magnetic beads. Isolated immune cells were then taken for downstream applications.

### Flow Cytometry

Brian immune cells and PBMC samples were resuspended in 200µL of cold 1XPBS and transferred into clear, v-bottom 96 well plate. Samples pelleted via centrifugation at 300xg for 5 minutes at 4°C and supernatant removed. Samples were washed once in 200µL of 1XPBS, pelleted via centrifugation (300xg, 5 minutes, 4°C), and resuspended in 50µL of antibody cocktail. Cells were incubated in the dark with antibody cocktail for 20 minutes at 4°C. After antibody incubation cells were pelleted and washed, 3 times, with Facs buffer. Cells were then fixed in 50µL of 1% PFA for 30 minutes at 4°C in the dark. After fixation cells were pelleted and washed twice in FACS buffer. For compensation controls, 1µL of antibody was added to one drop of reactive and negative AbC™ Total Antibody Compensation Beads, while 0.5µL of Live/Dead antibody was resuspended in one drop of reactive and negative ArC Amine Reactive Compensation Beads for live/dead compensation control. Samples and comps were resuspended in 200µL of FACS buffer and transferred to glass tubes which were topped off with another 100µl of FACS buffer. Samples were analyzed on the BD FACS symphony A3 until 100,000 single-cell live events were collected or until the sample ran dry. Brain immune cell and PBMC samples that did not capture at least 70,000 events or PBMC samples that did not capture over 50,000 live CD45+ events were excluded. Lasers were set with spherotech beads values determined by initial complete compensation conditions. Compensation controls were performed for each experiment and comps calculated prior to running samples. Samples from each treatment group were present in each run. FCS files were analyzed via Flowjo software version 10. Gates were set using Fluorescent minus one controls (FMOCs) (**Supplemental Figure 3**) and samples were normalized within sex between runs.

### RNA Extraction for Fresh Samples

Bench areas were cleaned with RNase Zap prior to RNA extraction. Brain immune cell isolation and PBMCs were lysed with 350µL BME and RLT lysis buffer (20µL of BME per 1mL RTL buffer) and processed for RNA extraction via RNeasy Kit (Qiagen) according to the manufacturer’s protocol. Briefly, lysed solution was transferred to Qiashredder tubes for complete homogenization. Sample flow-through was then mixed with 350µlL of 70% ethanol and transferred to RNAeasy spin columns. Samples were centrifuged for 15 seconds at 9391xg at room temperature. Flow-through was discarded and nucleic acids trapped in the RNAeasy membrane were washed with 700µL of RW1 buffer and 500µL of RPE buffer twice, centrifuging for 15 seconds at 9391xg at room temperature and discarding flow-through after each wash. Samples were then centrifuged for 2 minutes at full speed to dry before eluting samples with 20µL of RNAse free water in a 1 minute spin at 9391xg. SI

### NanoString

The NanoString nCounter® Analysis System (NanoString Technologies, Seattle, USA) was employed to perform the Mouse Immunology Panel nCounter® assay (CAT# xt_pgx_MmV1_Immunology_CSO). This assay utilized a panel consisting of 549 target genes with 14 internal reference genes and an additional set of 9 custom spike-in genes for RGS10, GPNMB, GRN, Histone 3, Histone Deacetylase 3, LRRK2, iNOS, P65 NFKB, and SIP. Brain immune cell and PBMC RNA samples were loaded and hybridized to the provided capture and reporter probes overnight at 65°C according to the manufacturer’s instructions. The samples were then added into the provided nCounter chip via use of the NanoString MAX/FLEX robot. Unhybridized capture and reporter probes were removed, and the hybridized mRNAs were immobilized for transcript count quantification on the nCounter digital analyzer. The data were then imported into the nSolver analysis software v4.0 for quality check and analysis according to the manufacturer’s instructions. Raw counts were normalized to the geometric mean of the positive and negative controls and count expression was calculated via nSolver Advanced Analysis v 2.0.134. Gene expression was only considered significant if *p* < 0.05 after adjustment using the Benjamini-Hochberg method for false discovery rates. Sample quality was determined via the DV200 index; and samples that did not meet the minimum RNA concentration threshold were excluded from analysis. The top 10 Gene pathways of differentially expressed genes were analyzed ShinyGO 0.77 set to mouse species and run through KEGG Pathway database. Functional sub-categories were identified within KEGG pathways diagrams that had consistent directionality of the DEGs.

### Statistical Analyses

Analyses for human data from the PPMI study were conducted in R studio.4.4.1 For the primary analysis of RGS10 and patient group, we used a 1-way ANCOVA with 3 levels: control, prodromal, and PD. In this analysis we covaried for sex and age. Furthermore, Pearson’s correlation test was used to compare RGS10 levels to age and UPDRS score in PD patients. Lastly, levels of RGS10 between males and females in each cohort were analyzed using type II ANOVA with Bonferroni correction for multiple comparisons. Analyses for mouse data were performed with Graph-Pad Prism 9. Group differences were analyzed using ordinary two-way ANOVA corrected for multiple comparisons with Tukey post-hoc test. Samples that are statistically different from each other do not share the same letter. *p* values ≤ 0.05 were considered statistically significant.

## Results

### RGS10 Protein levels are Reduced in the CSF of Individuals with PD Relative to Matched Controls

We utilized the proteomic dataset from project 151 of the MJFF PPMI study to assess RGS10 levels in the CSF of healthy controls (n=176), prodromal PD (n=345), and individuals with PD (n=607) (**Figure 1A**). Correcting for age and sex, we found a significant main effect of patient group (COHORT) on levels of RGS10 in the CSF (**Supplemental Figure 1A**), with post-hoc analysis identifying a significant decrease in RGS10 levels of PD patients compared to healthy controls and prodromal individuals with no difference between prodromal individuals and healthy controls (**Figure 1B**). Moreover, both covariates (age and sex) held large portions of the variance and were significant in their ability to predict RGS10 levels in the CSF (**Supplemental Figure 1A**). Therefore, we wanted to assess the relationship between RGS10 levels in the CSF and age using linear regression. In both healthy control and PD cohorts we find a significant but very weak negative correlation between RGS10 and age (**Figure 1C**). Additionally, males with PD have significantly less RGS10 in the CSF as compared to prodromal females or females with PD (**Figure 1D**). Lastly, as we saw a significant difference in the RGS10 levels between PD patients, we wanted to assess whether RGS10 levels have a relationship to disease progression. Using a linear regression, we found there was not a significant correlation between RGS10 levels and disease progression scored by total UPDRS scores (**Figure 1E**).

**Figure 1:**
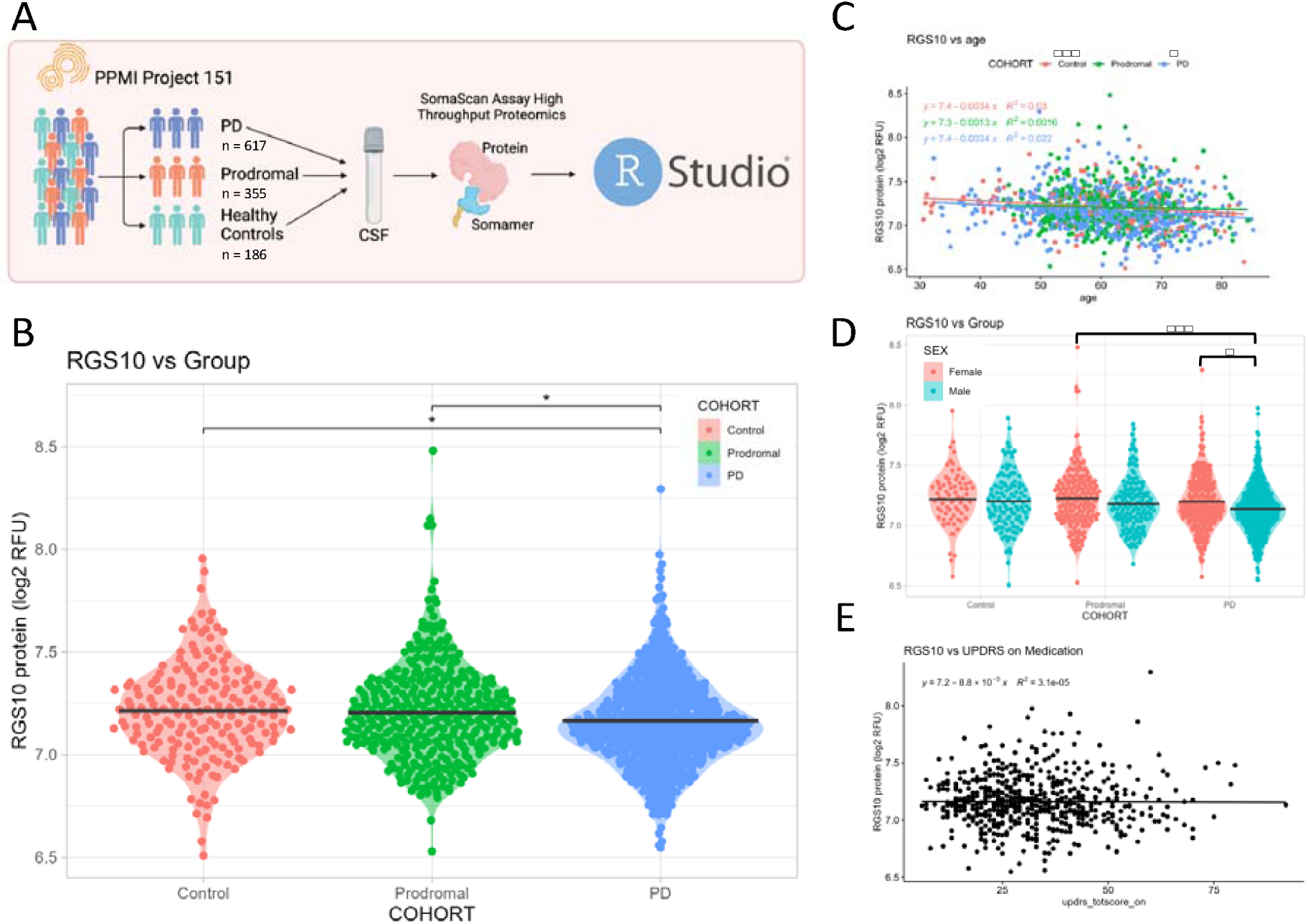
Individuals with PD display reduced protein levels of RGS10 in the CSF compared to healthy controls and prodromal individuals. A) Schematic of the study work flow. B) Log transformed quantification of relative fluorescence units (RFU) of RGS10 in the CSF of healthy controls, prodromal individuals, and individuals with PD. Statistical significance was calculated using 1-way ANCOVA with 3 levels: control, prodromal, and PD, covaried for sex and age, followed by Tukey’s HSD to correct for multiple comparisons. C) Log transformed quantification RFU of RGS10 across age in healthy controls, prodromal individuals, and individuals with PD. Statistical significance was calculated using a correlational test. D) Log transformed quantification RFU of RGS10 between sexes. Statistical significance was calculated using a 2-way Anova to compare RGS10 levels between male and females in each cohort. E) Log transformed quantification RFU of RGS10 across total UPDRS score in Parkinson’s Patients on medication. Statistical significance was calculated using a correlational test. *p < 0.05, ***p < 0.001.

As circulating, and likely immune cell populations in the CSF, suffer from a lack of RGS10 in individuals with PD[39], we aimed to investigate how loss of RGS10 impacts the inflammatory response in circulating and CNS-associated immune cells. Here, we investigated the immunoregulatory capacity of RGS10 under chronic systemic inflammatory conditions, which have been reported by multiple groups in individuals with PD and appear to be associated with the development of PD[9, 19, 20]. To do this we induced CSI in male and female C57BL6/J (B6) and RGS10 knock out (KO) animals. We isolated PBMCs and CNS-associated immune cells 24h after the completion of the CSI paradigm, thereby ensuring the wash out of acute inflammatory processes. Samples were subjected to comprehensive phenotyping of the innate and adaptive immune system via flow cytometry.

### RGS10 and CSI Regulate the Frequency and Functional Dynamics of Circulating Natural Killer Cells and Antigen-Presenting Cells in Mice

Considering, that loss of RGS10 promotes hyperinflammatory responses and dysregulated immune cell recruitment[22, 26, 28, 29, 32–34, 37, 38, 44], we assessed whether RGS10 mediates the frequency and functionality of innate immune cells exposed to LPS-induced CSI. Within the innate immune system, we found that LPS-induced CSI and RGS10 alter the frequency of patrolling natural killer (NK) cells, dendritic cells, and monocytes in the blood (**Figure 2A-C, F-H**). Specifically, we find a main effect of LPS-induced CSI and RGS10 on the frequency of NK cells, where KO animals have less NK cells overall but experience a significant increase in NK cell frequency with exposure to LPS-induced CSI, whereas B6 animals do not see a significant increase in NK cells frequency after LPS-induced CSI (**Figure 2A, F**). LPS-induced CSI also induced a significant increase in the frequency of dendritic cells compared to saline controls with RGS10 deficiency exacerbating this increase in male mice (**Figure 2B, G**). Lastly, we find that the percentage of circulating monocytes decreased in female (but not male) mice exposed to LPS-induced CSI (**Figure 2C, H**).

**Figure 2:**
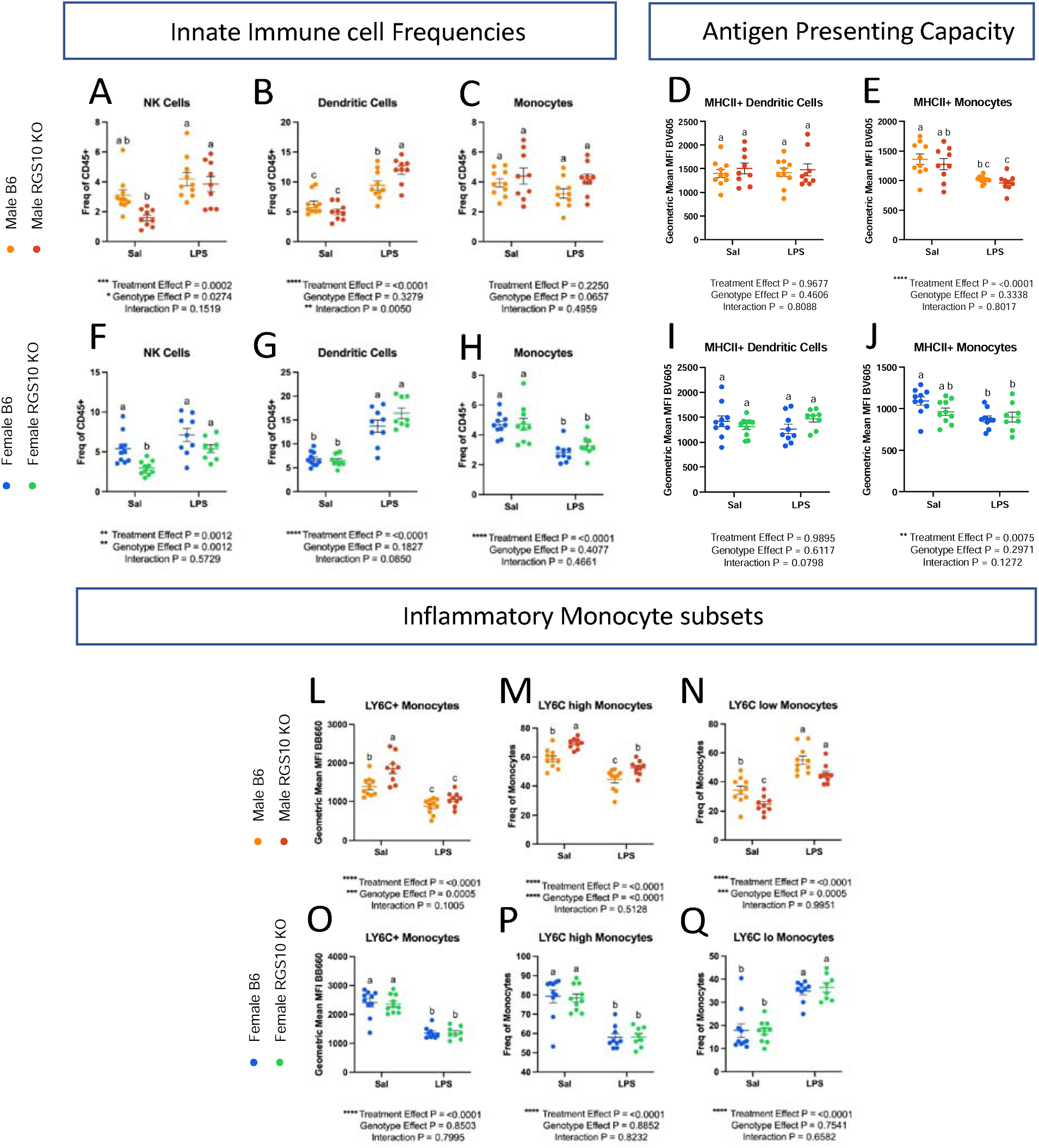
RGS10 and Chronic Systemic Inflammation (CSI) synergize to regulate the frequency and functional dynamics of natural killer cells and antigen-presenting cells in peripheral circulation. Male and female mouse PBMCs were immunophenotyped via flow cytometry. A & F) Frequency of natural killer cell out of total CD45+ cells. B & G) Frequency of dendritic cells out of total CD45+ cells. C &H) Frequency of monocytes out of total CD45+ cells. D & I) Mean fluorescent intensity (MFI) of MHCII on Dendritic cells. E & J) MFI of MHCII on monocytes. L & O) MFI of LY6C (BB660) on monocytes. M & P) Frequency of LY6C-high monocytes out of total monocytes. N & Q) Frequency of LY6C-low monocytes out of total monocytes. Samples that are statistically different do not share the same letter. Significance for main effects is as follows: *p < 0.05, **p < 0.01, ***p < 0.001, ****p < 0.0001.

To investigate functional changes within the antigen-presenting cell populations that were different between B6 and KO in the blood, we assessed MHCII membrane expression as a measure of antigen-presenting capacity and LY6C membrane expression as a marker of inflammatory monocytes. Despite a significant increase in the frequency of dendritic cells with CSI, we found that MHCII membrane expression on dendritic cells did not differ with LPS-induced CSI or RGS10 deficiency (**Figure 2D, I**). However, MHCII membrane expression was decreased on monocytes exposed to LPS-induced CSI (**Figure 2E, J**). Post-hoc analysis revealed deficits in MHCII membrane expression in male KO saline controls that was not present in female KO mice (**Figure 2E, J**). We also observed sex-specific differences as male, but not female, KO monocytes exhibited augmented inflammatory profiles (**Figure 2L-Q**). Specifically, membrane expression of LY6C on monocytes was markedly increased in KO male mice compared to B6 mice at baseline (**Figure 2L**). Moreover, RGS10-deficient male mice displayed a larger portion of LY6C-high monocytes (and inversely a lower proportion of LY6C-low) compared to B6 mice (**Figure 2M-N**). Interestingly, LPS-induced CSI decreased both the ratio and membrane expression of LY6C on monocytes regardless of sex (**Figure 2L-Q**). Overall, we found that RGS10 and LPS-induced CSI regulate the frequency and functional dynamics of NK cells and APCs in mouse PBMCs, with CSI increasing NK-cell and dendritic-cell frequencies in the blood, while dampening inflammatory phenotypes in antigen-presenting monocytes. Moreover, RGS10 deficiency exacerbated CSI-induced increases in dendritic cells and was associated with higher proportions of inflammatory monocytes, predominantly in male mice.

### RGS10 and CSI Regulate the Frequency of Circulating B and CD4+T cells in Mice

In light of the alterations in circulating APC frequency and capacity as a result of LPS-induced CSI and RGS10 deficiency described above, we also examined lymphocyte population dynamics in the circulation. Here, we found that LPS-induced CSI and RGS10 induce sex-specific alterations on B-cell frequency, while RGS10 regulates CD4+ helper T-cell population frequencies (**Figure 3**). Specifically, there is a main effect of CSI on B-cell frequency, with a significant reduction in B cells of B6 males but not KO males exposed to LPS-induced CSI (**Figure 3A, C**). The opposite is true in females, evinced by a significant reduction in B cells in KO females but not B6 females exposed to LPS-induced CSI (**Figure 3C**).

**Figure 3:**
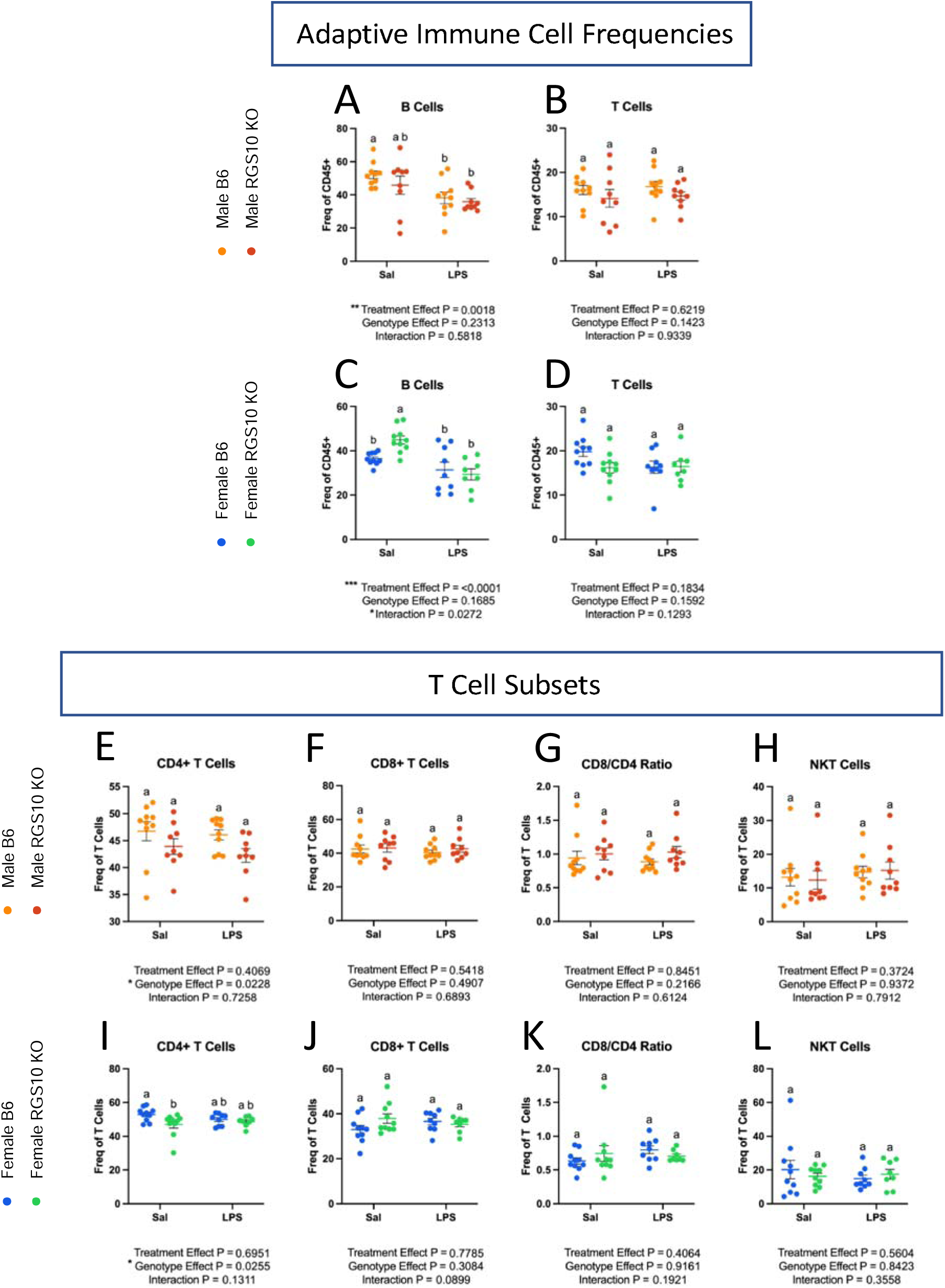
RGS10 and Chronic Systemic Inflammation (CSI) regulate the frequency of B and CD4+ T cells in circulation. Male and female mice PBMCs were immunophenotyped via flow cytometry. A & C) Frequency of B cells out of CD45+ cells. B & D) Frequency of T cells out of total CD45+ cells. Frequency of CD4+ cells out of total T cells in males (E) and females (I). Frequency of CD8+ cells out of total T cells in males (F) and females (J). CD8:CD4 T cell ratio in males (G) and females (K). Frequency of NK1.1+ cells out of total T cells in males (H) and females (L). Samples that are statistically different do not share the same letter. Significance for main effects is as follows: *p < 0.05, **p < 0.01, ***p < 0.001, ****p < 0.0001.

Overall, neither RGS10 deficiency nor LPD-induced CSI impacted the frequency of circulating T cells (**Figure 3B, D**). However, upon further evaluation of T-cell subsets, we found a main effect of RGS10 on CD4+ helper T cells (**Figure 3E, I**). In females, we also detected a significant decrease in CD4+ T cells in RGS10-deficient animals compared to B6 counterparts at baseline that disappears with LPS-induced CSI (**Figure 3I**). The frequency of CD8+ cytotoxic T cells and natural killer T (NKT) cells in the blood did not change as a result of CSI or RGS10 deficiency (**Figure 3F-H, J-L**). Interestingly, membrane expression of the primary markers for B cells (CD19), T cells (CD3), helper T cells (CD4), and cytotoxic T cells (CD8) were also decreased with RGS10 deficiency, primarily in males (**Supplemental Figure 7**). Taken together, these data indicate that LPS-induced CSI increases the frequency of patrolling cells within the circulation while decreasing the inflammatory and antigen-presenting capacity of monocytes. Additionally, CSI diminishes the frequency of B cells in the circulation. On the other hand, RGS10 deficiency exacerbates CSI-induced increases in dendritic cell populations and enhances the inflammatory status of monocytes in males, while reducing the frequency of CD4+ T cells in the circulation.

### RGS10 Deficiency Synergizes with CSI to Induce Innate and Adaptive Immune System Dysregulation in Male Mouse PBMCs

Next, we utilized a targeted transcriptomic assay of immune pathways enabled by Nanostring Technology to better understand the activation status and cellular pathways altered in RGS10-deficient immune cells exposed to LPS-induced CSI. To do this, we extracted RNA from PBMCs collected from a separate cohort of animals that were exposed to the same CSI paradigm. This separate cohort consisted of male mice based on the large proportion of the RGS10-dependent changes that we observed in males only.

Here we found that under baseline, non-stimulated conditions RGS10 deficiency did not induce any differentially expressed genes (DEGs) in PBMCs compared to B6 saline controls, with the exception of RGS10 itself (**Figure 4A**). Moreover, LPS-induced CSI alone only induces 8 DEGs relative to B6 saline controls (**Figure 4B**). However, after exposure to LPS-induced CSI, we found 117 DEGs in RGS10-deficient PBMCs as compared to B6 saline controls (**Figure 4C**), indicating that RGS10 primarily mediates immune pathways upon stimulation, an observation which is also evident in the sizable induction of DEGs in KO exposed to LPS-induced CSI when compared to KO saline controls (**Figure 4B**). Interestingly, we find no differentially expressed genes between KO and B6 PBMCs that were both exposed to LPS-induced CSI (**Figure 4A**), suggesting that CSI induced similar changes in immune cell pathways. We can further appreciate the synergy of RGS10 deficiency and LPS-induced CSI in altering immune pathways by comparing the similarity of DEGs between each group when compared to baseline. Here, we find that all 8 DEGs induced by LPS-induced CSI alone are shared with the KO group that was also exposed to LPS-induced CSI, furthermore 108 unique DEGs appeared in the KO group upon exposure to LPS-induced CSI (**Figure 4D**). Importantly, we also confirmed the absence of RGS10 in both of our KO groups (**Figure 4D**).

**Figure 4:**
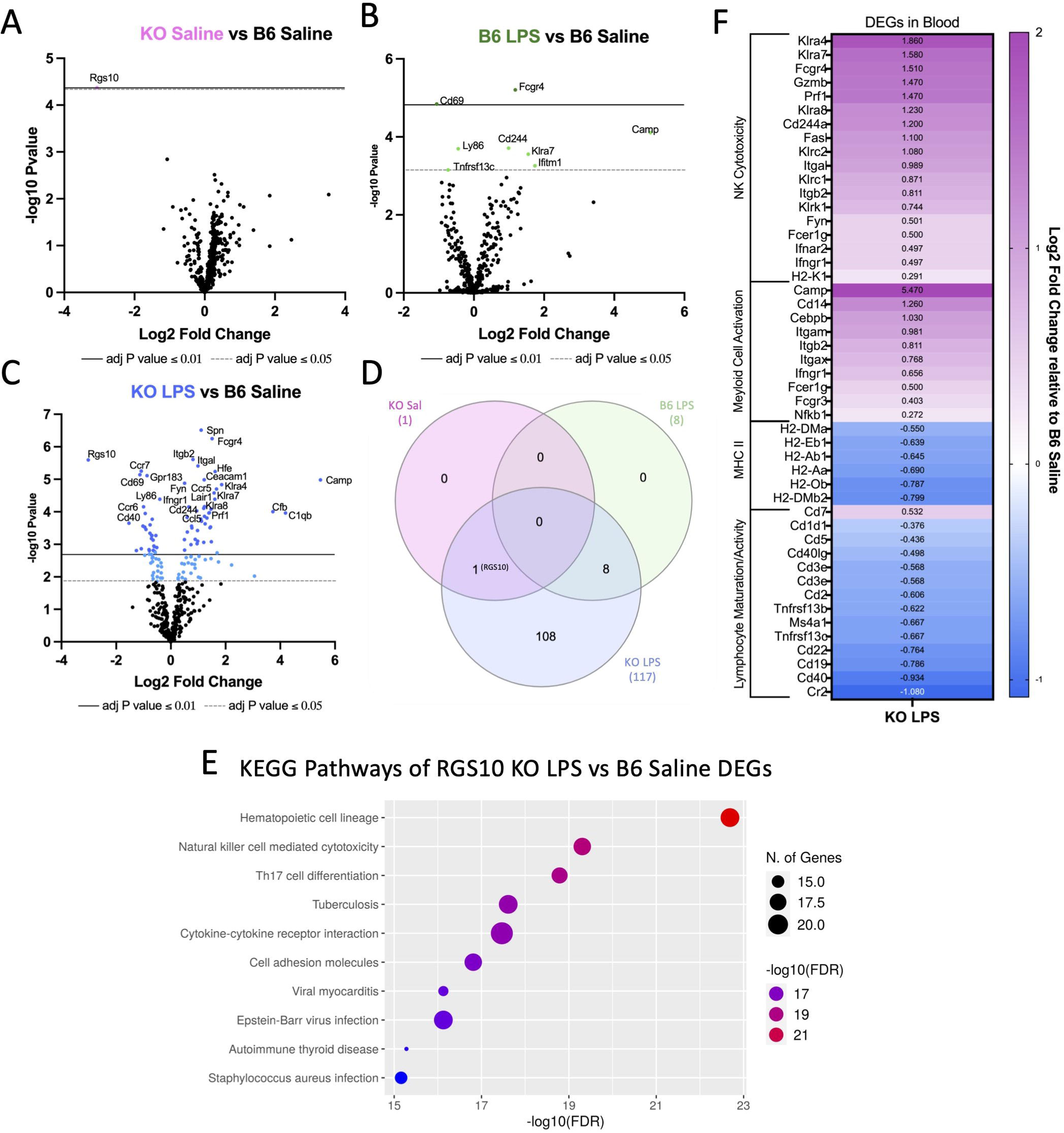
RGS10 deficiency synergizes with Chronic Systemic Inflammation (CSI) to induce innate and adaptive immune system dysregulation in mouse PBMCs. RNA from mouse PBMCs were run on the NanoString nCounter® immune panel. A) Volcano plot of differentially expressed genes (DEGs) of RGS10 KO Saline vs B6 Saline. B) Volcano plot of DEGs of B6 LPS vs B6 Saline. C) Volcano plot of DEGs of RGS10 KO LPS vs B6 Saline. D) Venn diagram of the number of shared and unique DEGs from each group compared to B6 Saline. E) Dotplot of 10 most significant KEGG pathways the DEGs from RGS10 KO LPS vs B6 Saline comparison are involved in. F) Heatmap of the fold change of RGS10 KO LPS and B6 LPS DEGs relative to B6 saline. All genes listed are significantly different in the RGS10 KO LPS group compared to B6 Saline (BH adjusted P < 0.05). Genes are grouped by associated KEGG pathways.

To analyze the gene pathways that were changing as a result of RGS10 deficiency and CSI, we ran all 117 DEGs identified in the KO group exposed to CSI through a KEGG analysis. The KEGG pathway analysis showed that DEGs of RGS10-deficient PBMCs exposed to LPS-induced CSI were involved in immune cell maturation/differentiation, cytokine-cytokine receptor interactions, cell-adhesion molecules, NK cell-mediated cytotoxicity, autoimmune thyroid disease, and a multitude of gene pathways in infection (**Figure 4E**). As these pathways included both up and downregulated DEGs, we performed sub-analyses in which we identified 4 functional sub-categories within KEGG pathways that had consistent directionality of the DEGs (**Figure 4F**). We found a significant upregulation of NK cell cytotoxicity and myeloid cell activation along with a significant downregulation in MHCII expression and lymphocyte maturation/activation in CSI-exposed RGS10-deficient PBMCs as compared to B6 saline controls (**Figure 4F**).

### RGS10 and CSI Regulate the Frequency of Innate Immune Cells in and around the Brain and Synergistically Influence their Activation

As immune cell phenotypes are influenced by origin and environmental context [45, 46], immune cells within the circulation may not reflect immune-cell populations in and around the brain. Therefore, we isolated brain immune cells to assess whether CNS-associated immune cells exhibit similar changes to those seen in circulation under conditions of RGS10 deficiency and LPS-induced CSI. Importantly, animals were not perfused prior to isolating brain immune cells, as we did not want to restrict our assessment of immune cells to that solely of the brain parenchyma as both meningeal and peri-vascular spaces can contribute to the induction of neuroinflammation[17]. Furthermore, we reported above that PD patients have less RGS10 in their CSF compared to healthy controls and prodromal individuals, highlighting the potential role of RGS10 in brain-adjacent compartments.

We investigated the frequency of CD45+ cells out of total live cells to determine the proportion of immune cells in and around the brain. Overall, we found a main effect of genotype on the proportion of CD45+ cells, demonstrating a reduction in the frequency of CD45+ cells in KO mice in both males and females (**Figure 5A-B**). However, upon post-hoc analysis there were no significant differences in the frequency of CD45+ cells in and around the brain with or without RGS10 deficiency or LPS-induced CSI in males (**Figure 5A**). Interestingly, however, we see a main effect of CSI treatment in females, that with post-hoc analysis reveals a significant increase in the frequency of CD45+ cells with CSI in B6 mice, but not in KO mice (**Figure 5B**). Moreover, we assessed membrane expression of CD45 as an indicator of immune activation in CNS-associated immune cells[47].CD45 may also reflect the proportion of peripheral immune cells present in the brain as peripheral immune cells express CD45 to a greater extent than CNS-resident immune cells like microglia[47]. Here we observed a main effect of LPS-induced CSI on CD45 membrane expression, with post-hoc analysis revealing that LPS-induced CSI, in males, significantly increased CD45 expression only in KO animals compared to saline controls, and not in B6 animals (**Figure 5C**). Conversely, we found a significant increase in CD45 expression due to LPS-induced CSI in females in both KO and B6 animals (**Figure 5D**).

**Figure 5:**
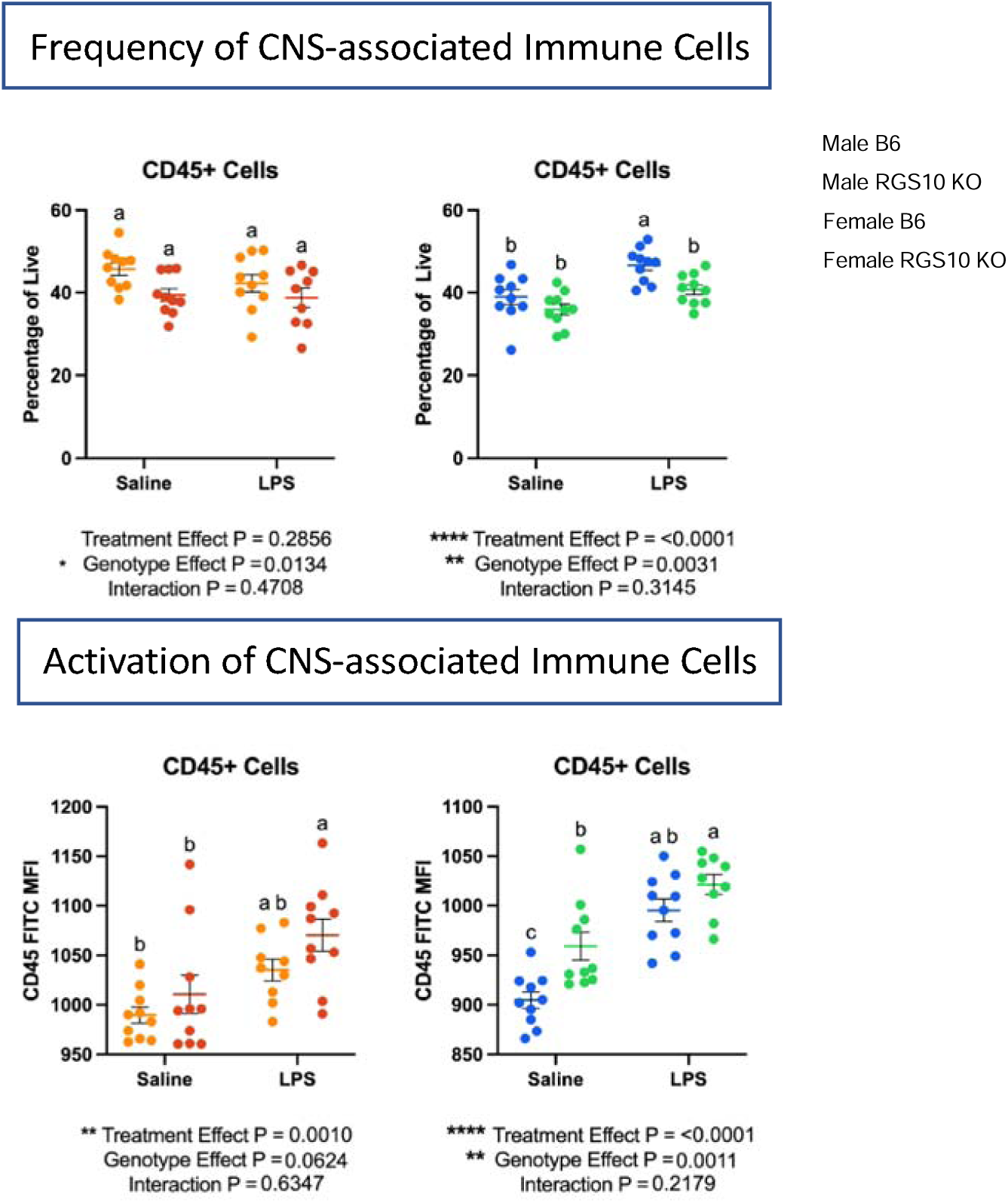
RGS10 and Chronic Systemic Inflammation (CSI) regulate immune cell frequency and activation in the CNS. Mouse PBMCs were immunophenotyped via flow cytometry. Frequency of CD45+ cells out of all live cells in males (A) and females (B). MFI of CD45 out of CD45+ immune cells in males (C) and females (D). Samples that are statistically different do not share the same letter. Significance for main effects is as follows: *p < 0.05, **p < 0.01, ***p < 0.001, ****p < 0.0001.

Furthermore, we also observed a significant increase in CD45 expression in KO females at baseline compared to B6 females (**Figure 5D**). Overall, these data indicate that RGS10 and CSI may regulate the frequency and activation of CNS-associated immune cells. Next, we investigated specific innate immune cell responses mediated by CSI and RGS10 in the CNS. Importantly, we assessed immune cell populations out of total CD45+ cells to normalize any differences between groups in immune cells present in the CNS. Overall, we found that LPS-induced CSI and RGS10 altered innate immune cell frequencies within the CNS. Most strikingly, we observed a marked increase in the frequency of dendritic cells in and around the brain of KO animals regardless of treatment or sex (**Figure 6A,E**). Moreover, we do not observe the same induction of NK cells in CNS-associated immune cells as we do in the circulation, which may be a product of the limited amount of NK cells in and around the brain (**Figure 6B,F**). Specifically, we observe a main effect of LPS-induced CSI on NK cells in and around the brain in males but find no significant differences in the frequency of NK cells between treatment groups regardless of sex (**Figure 6B,F**). Additionally, we found no differences in the frequency of CNS-associated monocytes in males but detected an increase in monocytes in females with LPS-induced CSI in B6 but not KO animals (**Figure 6C, G**). Lastly, we demonstrated a significant reduction in the frequency of microglia in KO animals compared to B6 saline controls in both sexes (**Figure 6D, H**), indicating an increase in frequency of peripheral immune cells out of the total immune cells present.

**Figure 6:**
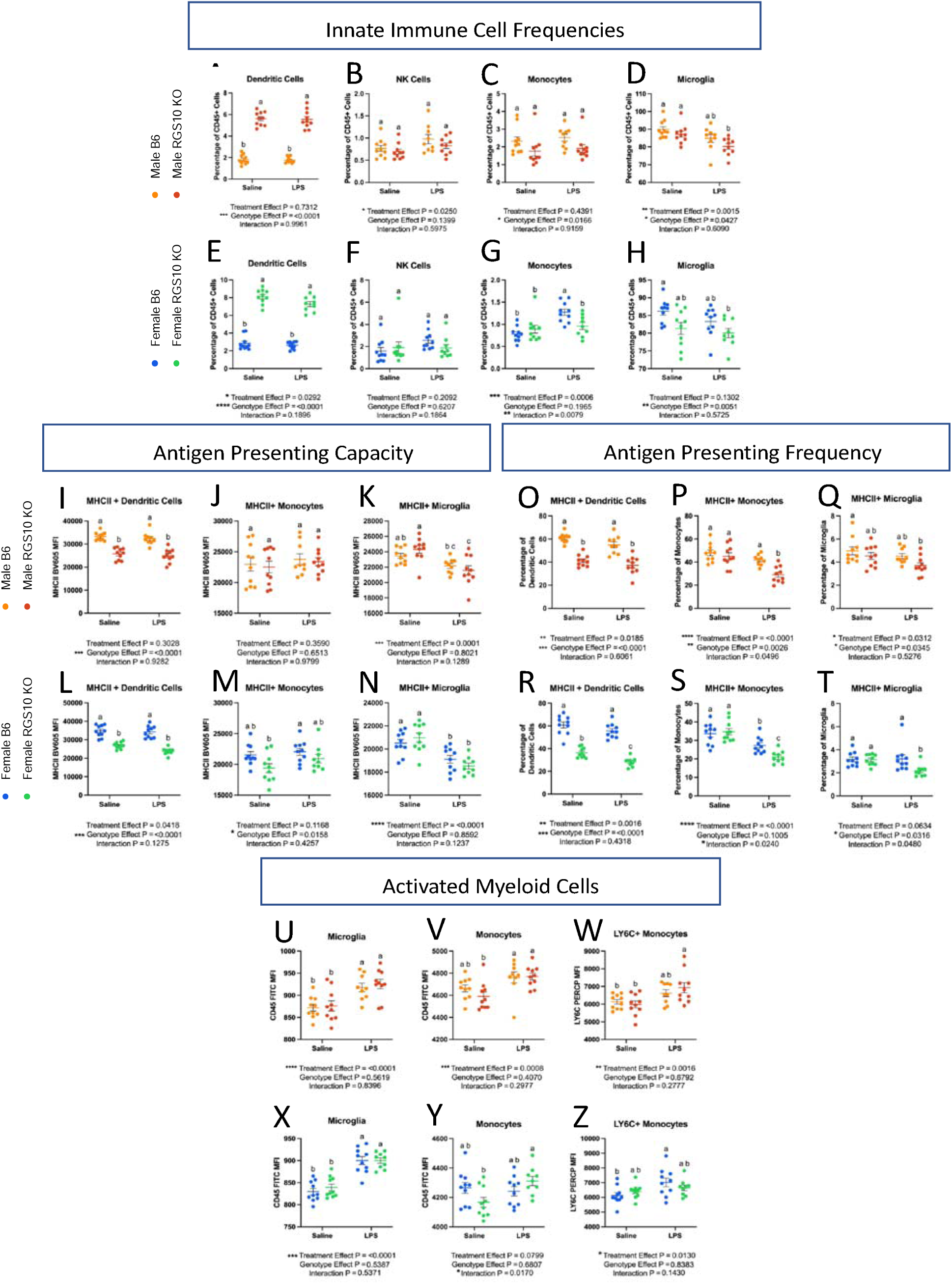
RGS10 and Chronic Systemic Inflammation (CSI) impair capacity for antigen presentation in brain-associated immune cells and induce myeloid cell activation. Male and female mouse CNS-associated immune cells were immunophenotyped via flow cytometry. A & E) Frequency of dendritic cells out of CD45+ cells. B & F) Frequency of natural killer cell out of CD45+ cells. I & L) Mean fluorescent intensity (MFI) of MHCII on dendritic cells. J & M) MFI of MHCII on monocytes. K & N) MFI of MHCII on microglia. O & R) Frequency of MHCII+ dendritic cells out of total dendritic cells. P &S) Frequency MHCII+ monocytes out of total monocytes. Q &T) Frequency MHCII+ microglia out of total microglia. U & X) MFI of CD45 on microglia. V & Y) MFI of CD45 on monocytes. W & Z) MFI of LY6C on monocytes. Significance for main effects is as follows: *p < 0.05, **p < 0.01, ***p < 0.001, ****p < 0.0001.

Unlike the blood, we found a significant decrease in MHCII membrane expression in RGS10-deficient dendritic cells in and around the brain (**Figure 6I,L**). Additionally, we see that the frequency of MHCII+ dendritic cells declined with RGS10 deficiency, and this was exacerbated in female KO mice when exposed to LPS-induced CSI (**Figure 6O,R**). Again, unlike the blood, we saw no differences in MHCII membrane expression in monocytes with CSI or KO compared to B6 saline controls (**Figure 6J, M**). However, the frequency of MHCII+ monocytes was significantly reduced with LPS-induced CSI in RGS10-deficient animals compared to all other groups in both males and females (**Figure 6P,S**). Moreover, microglia are also antigen-presenting cells, and we found a main effect of LPS-induced CSI on MHCII membrane expression on microglia along with a significant decrease in the frequency of MHCII+ microglia with LPS-induced CSI in KO mice compared to B6 saline controls, in the CNS (**Figure 6K,N,Q,T**). Overall, our data suggest a prominent decrease in antigen-presenting capacity in RGS10-deficient CNS-associated APCs exposed to CSI through reductions in membrane expression of MHCII and/or frequency of MHCII+ APCs.

Considering the overall activation of immune cells and increased presence of peripheral immune cells in the CNS with CSI and RGS10 deficiency we directly analyzed the inflammatory status of monocytes and microglia. Here, we observed a significant increase in CD45 membrane expression on microglia with LPS-induced CSI (**Figure 6U, X**). Additionally, we found a significant induction of CD45 expression on monocytes in and around the brain of RGS10-deficient animals with CSI but not in B6 animals (**Figure 6V, Y**). It should be noted, however, that there is no difference in CD45 expression level between B6 and KO monocytes after exposure to LPS-induced CSI, suggesting that KO monocytes may have slightly lower baseline levels of CD45. Lastly, looking at LY6C membrane expression, we observed increased LY6C expression on RGS10-deficient CNS-associated monocytes from male mice exposed to CSI as compared to saline controls. Conversely, LY6C membrane expression on CNS-associated monocytes was augmented with LPS-induced CSI in B6 females but not KO females (**Figure 6W, Z**). Collectively, these data reveal that myeloid cells in the CNS, and in particular monocytes, express robust induction of inflammatory markers after exposure to CSI and become activated in response to CSI with RGS10 deficiency, most prominently in males.

### RGS10 and CSI Mediate Reductions in the Lymphocyte Frequency while Enhancing Cytotoxic T Cell Populations in CNS-Associated Immune Cells

We then examined lymphocyte dynamics in the CNS. We found a reduction in the frequency of B cells (**Figure 7A, C**). In males, all groups displayed a significant reduction in the frequency of B cells compared to B6 saline controls (**Figure 7A**). In females, only B cells from KO mice exposed to LPS-induced CSI were significantly reduced compared to B6 saline controls (**Figure 7C**). Interestingly, we found a main effect of CSI on the frequency of T cells in the CNS (**Figure 7B, D**). When we broke down the T cells into their respective subsets, we found a main effect of LPS-induced CSI on each T-cell subset (**Figure 7E-L**). Specifically, there is a decreased frequency of CD4+ T cells in CSI-exposed animals in males. Importantly, we also observed that RGS10-deficient animals exhibited a lower frequency of CD4+ T cells at baseline, but we did not see any group differences in females (**Figure 7E, I**). Moreover, we found a significant increase in the frequency of CD8+ T cells with LPS-induced CSI; however, in males, this increase is only significant with RGS10 deficiency (**Figure 7F, J**). These changes in the frequency of CD4+ and CD8+ T cells were reflected in a significant increase in the ratio of CD8+ to CD4+ cells in KO males exposed to LPS-induced CSI (**Figure 7G**), indicating a shift to more cytotoxic T cell populations in RGS10-deficient animals exposed to LPS-induced CSI, most prominently in males. Expanding upon our analysis of cytotoxic T cell subsets we also found a significant increase in the frequency of NKT cells with LPS-induced CSI, however, in males, this increase was only significant with RGS10 deficiency (**Figure 7H, L**). Therefore, in CNS-associated immune cells we observed a larger frequency of peripheral immune cells in and around the brain of KO animals exposed to LPS-induced CSI, a massive increase in dendritic cells with RGS10 deficiency which may possibly act as a compensatory mechanism for the global impairment of MHCII expressing APCs in the CNS with RGS10 deficiency and LPS-induced CSI, as well as the enhancement of inflammatory myeloid cells and cytotoxic T cells subsets with CSI and KO.

**Figure 7:**
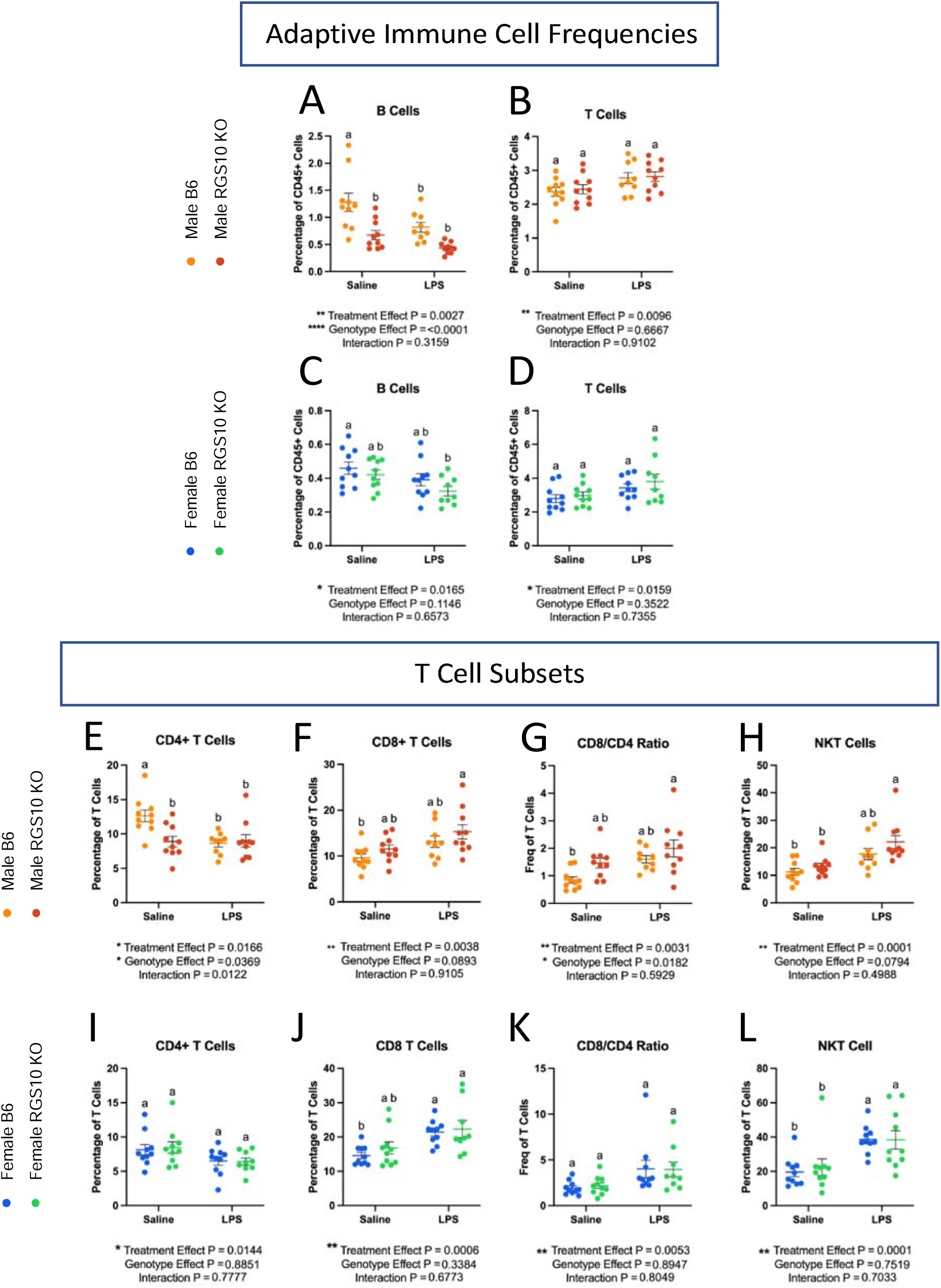
RGS10 and Chronic Systemic Inflammation (CSI) mediate reductions in the lymphocyte frequency while enhancing cytotoxic T cell populations in CNS-associated immune cells. Mouse CNS-associated immune cells were immunophenotyped via flow cytometry. A & C) Frequency of B cells out of CD45+ cells. B & D) Frequency of T cells out of CD45+ cells. Frequency of CD4+ cells out of total T cells in males (E) and females (I). Frequency of CD8+ cells out of total T cells in males (F) and females (J). Frequency of CD8/CD4 ratio out of total T cell in males (G) and females (K). Frequency of NK1.1+ cells out of total T cells in males (H) and females (L). Significance for main effects is as follows: *p < 0.05, **p < 0.01, ***p < 0.001, ****p < 0.0001.

### RGS10 Deficiency Synergizes with CSI to Activate Innate Immunity in CNS-Associated Immune Cells while impairing Lymphocyte Engagement

Assessing alterations in genes and gene pathways in CNS-associated immune cells, we observed an increase in the number of DEGs with exposure to LPS-induced CSI and RGS10 deficiency (**Figure 8A-D**). Interestingly, RGS10 uniquely regulated the differential expression of multiple genes within the CNS, unlike in the blood, including the downregulation of IL-6, IL-1β CCL6, CD11c, CD48, CD80, Fcgr4, Tnfsf12, Tnfsf13b, and H2-DMa, and upregulation of Mr1. Additionally, we found that CSI alone induced 3 DEGs (downregulating CCL24 and Cd164, and upregulating CXCL13) that were also induced with LPS -induced CSI in KO animals (**Figure 8A-D**). Moreover, RGS10 deficiency was sufficient to induce significant downregulation of Itgax, H2-DMa, and CD99 in CNS-associated immune cells when compared to B6 with both groups exposed to LPS-induced CSI (**Supplemental Figure 11C**), while there were no significant differences between KO and B6 groups exposed to LPS in the blood other than RGS10 (**Supplemental Figure 11D**). Running the 77 DEGs of CSI exposed KO animals through KEGG pathway analysis revealed that a large portion of DEGs were involved in infectious pathways or inflammation-driven diseases along with hematopoietic cell lineage, cytokine-cytokine receptor interactions, and IgA production (**Figure 8E**). Through our sub-analyses of the top 10 KEGG pathways distinguished in E, we identified 7 functional sub-categories with consistent directionality of the DEGs (**Figure 8F**). Here, we find that RGS10-deficient CNS-associated immune cells are enriched in TLR4/RIG1 pathways, complement genes, lymphocyte cell adhesion and migration genes, and IL-10 receptor genes (**Figure 8F**). Moreover, we find a bidirectional regulation of cytokines with enrichment in IL1a but a reduction in IL-6. Lastly, RGS10-deficient CNS-associated immune cells exposed to LPS-induced CSI exhibited significant reductions in chemokines related to myeloid trafficking and lymphocyte engagement through MHCII and costimulatory genes (**Figure 8F**). In summary, synergy of RGS10 deficiency and CSI regulates genes involved in immune cell trafficking, TLR4/RIG1 immune cell activation, cytokine and complement cascades, and lymphocyte engagement in CNS-associated immune cells.

**Figure 8:**
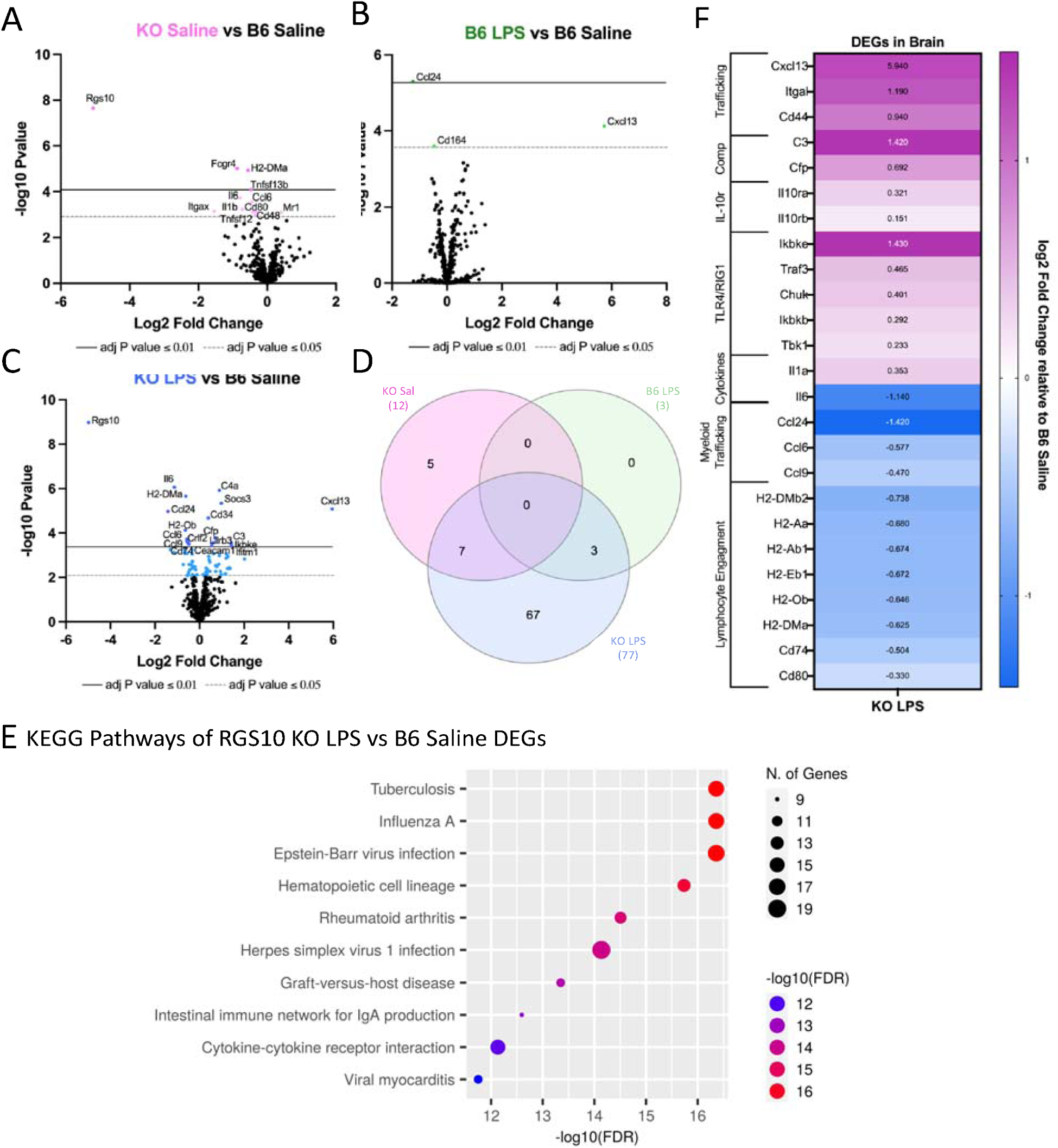
RGS10 deficiency synergizes with Chronic Systemic Inflammation (CSI) to activate innate immunity in CNS-associated immune cells while impairing lymphocyte engagement. RNA from mouse brain immune cells were run on the NanoString nCounter® immune panel. A) Volcano plot of differentially expressed genes (DEGs) of RGS10 KO Saline vs B6 Saline. B) Volcano plot of DEGs of B6 LPS vs B6 Saline. C) Volcano plot of DEGs of RGS10 KO LPS vs B6 Saline. D) Venn diagram of the number of shared and unique DEGs of each group compared to B6 Saline. E) Dot plot of 10 most significant KEGG pathways, the DEGs from RGS10 KO LPS vs B6 Saline comparison are involved in. F) Heatmap of the fold change of RGS10 KO LPS and B6 LPS DEGs relative to B6 saline. All genes listed are significantly different in the RGS10 KO LPS group compared to B6 Saline (BH adjusted P < 0.05). Genes are grouped by associated KEGG pathways.

## Discussion

As inflammation and immune dysregulation have been consistently shown to contribute to the pathophysiology of PD, it is pertinent to investigate changes in immunoregulatory protein levels of relevant immune cell populations such as CNS-resident and peripheral immune cells, to understand disease-relevant mechanistic changes and identify potential therapeutic targets[9]. Here, we demonstrate a decrease in RGS10, a critical homeostatic regulator of immune cells, in the CSF of individuals with PD compared to healthy controls and prodromal individuals. As RGS10 is most highly expressed in immune cells, decreased levels of RGS10 may indicate a reduction of RGS10 in immune cells in the CSF of individuals with PD. This suggests that peripheral immune cells capable of directly interacting with the CNS in individuals with PD may lack the ability to negatively regulate the inflammatory response at the same level of healthy controls and may engage hyperinflammatory responses or develop chronic inflammatory responses that endanger vulnerable neuronal populations[48]. Considering that RGS10 levels are also reduced in subsets of immune cells in the peripheral circulation in PD, we examined the role of RGS10 in regulating the immune response of circulating and CNS-associated immune cells to CSI, as CSI exemplifies inflammatory conditions associated with and present in PD. We demonstrate that RGS10 and CSI synergize to regulate myeloid cell activation, antigen-presenting capacity, and frequency of cytotoxic cell populations in both circulating and CNS-associated immune cells (**Figure 9**).

**Figure 9:**
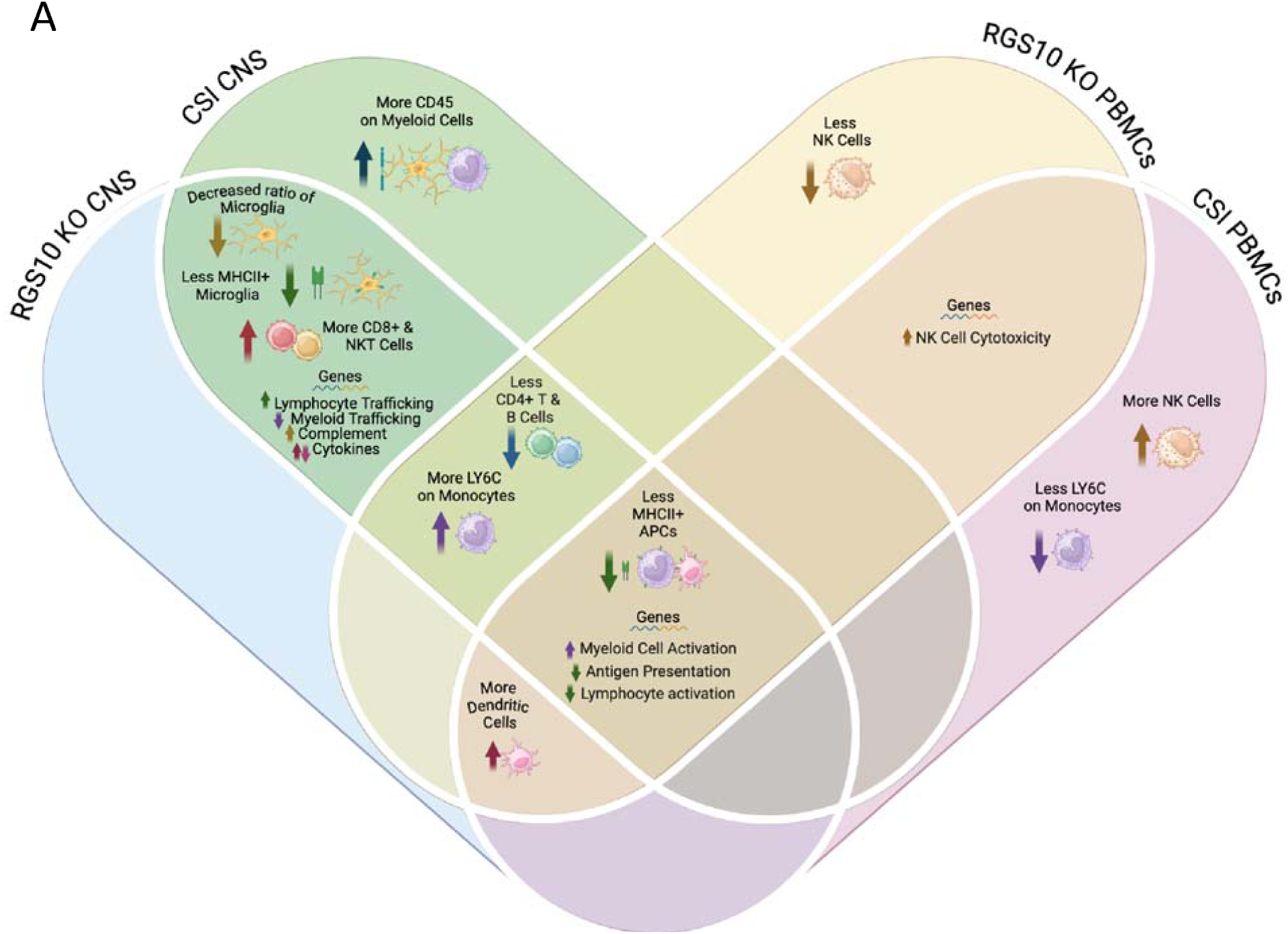
Summary of RGS10 and Chronic Systemic Inflammation (CSI)-mediated alterations in circulating and CNS-associated immune cells. A) Venn diagram of immune cell phenotypes identified in each group. Phenotypes that were present in both RGS10 KO, CSI, or the combination are listed in overlapping areas based on location in the CNS or blood. Common to both PBMCs and CNS-associated immune cells, RGS10 deficiency and CSI induce reductions in antigen presentation capacity, increased myeloid cell activation, and decreased lymphocyte engagement. Made with Biorender.com.

Notably, this is the largest study to investigate levels of RGS10 in individuals with PD. As such, this is the first study to assess RGS10 levels in a sufficiently large population to determine more than disease effect, identifying that RGS10 levels are modestly but significantly inversely correlated with age in controls and individuals with PD. This points to deficits in RGS10 being driven, in part, by aging, which is the largest risk factor for developing neurodegenerative diseases. Conversely, mouse studies have identified augmented levels of RGS10 in aged monocytes and granulocytes in the circulation, which may reflect a compensatory or tissue-specific phenomena in mice[49]. Regardless, the loss of RGS10 with age in the CSF is supported by the concept of immune dysregulation and inflammaging[50]. Specifically, it is important to highlight that inflammatory insults such as LPS induce downregulation of RGS10 via histone deacetylation[35]. Within our CSI paradigm, transcript levels of RGS10 were not significantly decreased in the PBMCs but were, however, in CNS-associated immune cells. We may have been able to appreciate significant changes in RGS10 transcript levels in CNS-associated immune cells due to the abundance of microglia which highly express RGS10 ubiquitously in the CNS[51, 52], whereas changes in the blood may not have been detected due to its heterogenous cellular makeup and/or rapid turnover of cells, indicating potential cell-specific alterations in RGS10 which could be assessed in follow-up experiments utilizing single-cell transcriptomics. Nevertheless, we present evidence that LPS-induced CSI and not just acute inflammatory stimuli can reduce RGS10 transcript levels in immune cells. Consequently, CSI-induced reductions in RGS10 may enhance the feed forward cycle of CSI.

Importantly, we are capturing immune frequencies 24 hours past the last injection of LPS to ensure that acute inflammatory responses have resolved[53]. As a result, one limitation of our analysis is that we miss the early influx of neutrophils and monocytes into the circulation for homing to the peritoneum, the site of injury[54]. However, this time scale allows us to investigate stable and more long-lasting alterations in the frequency of immune cells in the circulation. Moreover, due to the chronicity of the paradigm we would expect both APC activity and lymphocytic participation. Here, we demonstrate that LPS-induced CSI alone can increase the frequency of NK cells, dendritic cells, and patrolling LY6C-low monocytes. These cell types are specialized in surveillance and likely increased in response to persistent presence of LPS in the circulation. Specifically, LPS has been shown to induce NK cell proliferation as well as promote dendritic cell maturation and survival in vitro[55, 56]. LPS has also been shown to enhance LY6C expression on monocytes[57], however, we find that LPS-induced CSI increased the proportion of LY6C-low monocytes. High expression of LY6C is associated with a classical monocyte phenotype, which is characterized to be more pro-inflammatory in the blood; however, high expression of LY6C has been deemed a transient phenomenon as LY6C levels drop upon recruitment to lymphoid and non-lymphoid tissue[46]. Therefore, we may be capturing the conversion of LY6C-high monocytes to LY6C-low patrolling monocytes within the circulation with LPS-induced CSI.

Moreover, increased monocytes in the blood have been reported in chronic infection and chronic auto-inflammatory disorders[58]. In contrast, in our model we find no difference in monocyte frequency in males with LPS-induced CSI and a decrease in the frequency of monocytes with LPS-induced CSI in female animals. This difference may arise for a few different reasons. Firstly, as mentioned previously, we did not capture acute monocyte dynamics that may be more reflective of active infections rather than low-grade systemic inflammation. Secondly, monocytes can differentiate into tumor necrosis factor-and inducible nitric oxide synthase-producing dendritic cells (TIP-DCs), and our concurrent increase in dendritic cells with CSI may reflect increased monocyte to DC differentiation[46]. Lastly, in vitro work has demonstrated that chronic LPS-induced reductions in MHCII couple with increases in PD-L1 expression, demarcating exhausted monocytes[59]. Considering that LPS-induced CSI also decreased MHCII membrane expression on monocytes in circulation, our decreased frequency of monocytes may be related to monocyte exhaustion[59]. Furthermore, the frequency of B cells in the peripheral circulation were decreased as a result of LPS-induced CSI. This reduction in B cells may be due to decreased lymphopoiesis from hematopoietic output in favor of myelopoiesis under inflammatory conditions[60].

Notably, we find that RGS10-and CSI-mediated alterations in gene pathways corroborate and extend our flow cytometry results. Specifically, we find that even though KO animals have less NK cells overall, the increase in NK cells with LPS-induced CSI was only present in KO animals. Moreover, KO animals exposed to LPS-induced CSI, experienced a significant upregulation of genes involved in natural killer-cell cytotoxicity compared to B6 saline controls. Thus, we report herein a novel role of RGS10 in NK cell frequencies and activation which offers an exciting new line of investigation into the role of RGS10 in the regulation of the innate immune system. Additionally, upregulation of genes involved in myeloid activation are consistent with higher LY6C expression in RGS10-deficient monocytes and the capacity of RGS10 to regulate the inflammatory response of myeloid cells[22, 29, 32–34, 37]. Furthermore, the reduction in MHCII-related genes supports our observation of impaired antigen-presenting capacity in monocytes as a result of LPS-induced CSI. Correspondingly, we also find that LPS-induced CSI and RGS10 mediate the downregulation of genes involved in lymphocyte activation/maturation, along with RGS10 specific deficits in CD4+ T cell frequency in the circulation, consistent with previous studies[23, 49]. As this deficit in CD4+ T cells was also present in the CNS, this would suggest a systemic reduction in CD4+ T cells indicating that RGS10-deficient animals may have impaired CD4+ T cell engagement and expansion or enhanced CD4+ T cell death. Lastly, we find that downregulation of CD3 and CD19 is consistent with the reduction of their respective membrane expression on T and B cells with RGS10 deficiency. In brief, our data reveal a synergistic relationship with RGS10 deficiency and LPS-induced CSI that alters homeostasis to induce significant differentially-expressed genes denoting a shift to more cytotoxic and inflammatory innate immune populations that have impaired ability to talk to the adaptive immune system, thereby impairing its activation and potential resolution of the inflammatory response.

In the CNS, we do not observe the same NK cell phenotypes as in the blood, possibly due to the low number of NK cells found in the CNS. Moreover, unexpectedly and in contrast to our findings in blood, we find a significant increase in dendritic cells that is not mediated by LPS-induced CSI, but solely by RGS10 deficiency, revealing a novel role of RGS10 in dendritic cell biology within the CNS. Monocyte populations in the CNS, on the other hand, were similar to those in the blood in that we see no changes in frequency between groups for males. Interestingly though, we found a significant increase in the frequency of monocytes with LPS-induced CSI in females but only in WT B6 animals, not KO animals, supporting a role for RGS10 in sex and genotype effects on CNS-associated monocyte population dynamics. Consistent with the general reduction or lack of expansion of monocytes in RGS10-deficient animals, we find that genes for multiple chemokine ligands important for monocyte trafficking are downregulated in the CNS-associated immune cells of KO animals exposed to LPS-induced CSI. Regardless of immune cell frequency, we see amplified activation of immune cells in the CNS and specifically in myeloid populations, including microglia, with enhanced CD45 expression with LPS-induced CSI and LY6C expression on monocytes with RGS10 and LPS-induced CSI, consistent with increased LY6C on circulating monocytes. Moreover, genes related to myeloid activation, specifically, TLR4/RIG1 signaling and complement, were upregulated in CNS-associated immune cells with CSI of KO animals. Genes related to proinflammatory cytokines, however, were not regulated uniformly, with significant downregulation of IL-6 and upregulation of IL-1α. Interestingly, KO animals also displayed attenuated IL-6 increases in the CSF with aging as compared to B6 animals at baseline[49]. It would be pertinent to assess the protein level of these cytokines in the CNS to understand whether altered transcription in KO animals after CSI reflects concurrent protein levels or upstream regulation of adjusted protein requirements by the cell.

Moreover, we also observe reductions in antigen-presenting capacity in the CNS and the blood; however, all APC populations within the CNS, including microglia, experience decreased MHCII membrane expression or decreased frequency of MHCII+ cells with RGS10 deficiency and LPS-induced CSI. Dendritic cells displayed a significant genotype effect on reduced MHCII expression and or/MHCII+ frequency in the CNS. We hypothesize that the reduction in the frequency of CD4+ T cells with LPS-induced CSI and RGS10 deficiency may be a result of this decreased antigen-presenting capacity, especially considering that CD4+ T cells are preferentially expanded in bacterial infections which the presence of LPS would mimic[61]. This is further supported by the downregulation of genes involved in lymphocyte engagement in KO CNS-associated immune cells exposed to LPS-induced CSI. Furthermore, decreased frequencies of B cells with KO and LPS-induced CSI may also be related to the lack of CD4+ T cells that help to initiate the activation and proliferation of B cells[61]. Intriguingly, even though CD4+ T cell frequencies are reduced, we find an increase in the frequency of all CD3+ T cells in and around the brain with LPS-induced CSI. Moreover, genes that are important for immune cell adhesion and migration, particularly lymphocyte adhesion and migration, were upregulated in RGS10-deficient CNS-associated immune cells exposed to LPS-induced CSI. Correspondingly, KO animals displayed increased frequencies of cytotoxic T cell subsets (CD8+ and NKT cells) with CSI, which would be responsible for the overall increase in T cells and highlighting a shift towards cytotoxic peripheral immune cells in and around the brain. Interestingly, T cell chemotaxis was found to be inhibited, granting protection in and EAE model[38], indicating that RGS10 may differentially regulate immune cell trafficking in autoimmune versus chronic inflammatory conditions.

Overall, we found no significant differences in the total percentage of CNS-associated immune cells between groups in our males or our KO females, but did observe an increase in CD45+ cells with LPS-induced CSI in B6 female mice which reveal important sex and genotype specific effects on CNS-associated immune cell population dynamics. Interestingly, we see a bias for more CNS-associated phenotypes due to RGS10 deficiency and LPS-induced CSI than blood-associated phonotypes, indicating that CNS-associated immune populations may be more sensitive to loss of RGS10 under chronic inflammatory conditions which is consistent with the fact that RGS10 transcripts were significantly reduced in the CNS but not in the blood. Moreover, we were able to detect significant sex-specific differences with RGS10 in both humans and mice. Specifically, within PD, males have less RGS10 than females. This difference is interesting considering that men are at a higher risk for developing PD than women[62].

Previous studies using mouse models demonstrate that RGS10-deficient male mice develop inflammatory related phenotypes earlier and to a greater degree compared to their female counterparts[39]. Our study, highlights the sex-specific alterations in immune cell frequencies and activation with RGS10, where most of the phenotypes described are more prominent or only present in males, which is particularly evident in myeloid activation.

In summary, we demonstrate a reduction in RGS10 levels in the CSF of PD patients compared to healthy controls and prodromal individuals. Furthermore, we reveal significant alterations in myeloid cell activation, antigen-presenting capacity, and cytotoxic immune cells in the blood and CNS under CSI conditions when RGS10 is absent. Importantly, we cannot determine if this reduction of RGS10 in the CSF is specific to immune cells, as the proteomic data set lacks single cell resolution. This limitation could be mitigated by single-cell proteomics or mass cytometry analyses in future studies. Single-cell transcriptomic analysis and ATAC-sequencing of the CSF would also reveal if RGS10 transcripts were suppressed in PD patients through epigenetic regulation. Additionally, if longitudinal proteomic data from within subjects becomes available, re-running these analyses as repeated measures within subjects would allow us to better determine if RGS10 levels change with PD progression, advanced age, and/or years with disease.

This study examined RGS10 mediated immune functions in adult mice but not aged mice. To better understand RGS10 mediated immunity in the context of neurogenerative diseases, in future studies we plan to examine our CSI paradigm in the context of aging. Moreover, our assessment of immune cell populations through flow cytometry captures immune cells frequencies present at the time of harvest, and we are unable to determine immune cell recruitment and/or migration to or through our tissues of interest. Immune cell tracking and fate-mapping experiments would further inform the extent to which alterations in immune cell frequencies are a result of immune cell recruitment or local expansion. This would be especially informative to determine the extent of peripheral immune cell infiltration into the brain parenchyma. Accordingly, future studies should identify how RGS10 and CSI regulate specialized immune cell populations that reside in each CNS compartment (i.e. the parenchyma, the meninges, and the peri-vascular space) separately. Additionally, we utilized a targeted transcriptomic analysis that relies on reads of raw RNA counts and did not include any gene amplification therefore limiting the number of transcripts we are able to capture. Furthermore, as with the proteomic analysis above, our target transcriptomic analysis lacks single-cell resolution, and we are therefore unable to assign differentially expressed genes to particular cell types. Lastly, we observed significant impairments in antigen-presenting capacity as well as lymphocyte engagement due to KO and CSI which may indicate that RGS10-deficient immune cells are more primed for exhaustion under chronic inflammatory conditions and potentiate defective immune responses. Therefore, future studies should also examine the functional capacity of and status of immune cell exhaustion in RGS10-deficient immune cells exposed to CSI.

## Conclusions

Here we report a decrease in RGS10 in the CSF of individuals with PD compared to healthy controls and prodromal individuals. Additionally, we find that RGS10 levels are significantly predicted by age and sex but not disease progression as measured by the total UPDRS score. Moreover, we find that RGS10 deficiency synergizes with LPS-induced CSI to induce a bias towards inflammatory myeloid cells and cytotoxic cell populations, as well as a reduction in innate and adaptive crosstalk via MHCII in the circulation and in and around the brain most notably in males. These results, for the first time, highlight RGS10 as an important regulator of the systemic immune response to CSI and implicate RGS10 as a potential contributor to the development of immune dysregulation in PD.

## Abbreviations

RGS10: Regulator of G-protein signaling 10
PD: Parkinson’s disease
CSF: Cerebral spinal fluid
PPMI: Parkinson’s Progression Markers Initiative
CSI: Chronic systemic inflammation
CNS: Central nervous system
NFK-kB: Nuclear factor kappa-light-chain-enhancer of activated B cells
STIM2: Stromal interaction molecule 2
ROS: Reactive oxygen species
CIDs: Chronic inflammatory diseases
RA: Rheumatoid arthritis
IBD: Inflammatory bowel disease
IBS: Irritable bowel syndrome
RNA: Ribonucleic acid
PBMCs: Peripheral mononuclear cells
MJFF: Michael J Fox Foundation
IRB: International review board
DAT: Dopamine
RFU: Relative fluorescence units
pQTL: Protein quantitative trait loci
IACUC: Institutional Animal Care and Use Committee
B6: C57BL6/J
KO: RGS10 KO
LPS: Lipopolysaccharide
IP: Intraperitoneal
EDTA: Ethylenediaminetetraacetic acid
ACK: Ammonium chloride potassium
HBSS: Hank’s balanced salt solution
PBS: Phosphate buffered solution
D-PBS: Dulbecco’s Phosphate-Buffered Saline
GPNMB: Transmembrane glycoprotein nmb
GRN: Granulin
LRRK2: Leucine-rich repeat kinase 2
iNOS: Inducible nitric oxide synthase
SIP: Shingosine1-phosphate
ANCOVA: Analysis of covariance
ANOVA: Analysis of variance
NK: Natural Killer
MHCII: Major histocompatibility complex II
LY6C: Lymphocyte antigen 6 family member C
APC: Antigen presenting cell
DEG: Differentially expressed genes
IL-6: Interleukin 6
IL-1B: Interleukin 1 Beta
CCL6: Chemokine ligand 6
Fcgr4: FC receptor, IgG, low affinity IV
Tnfsf12: Tumor necrosis factor super family member 12
Tnffsf13b: Tumor necrosis factor super family member 13b
H2-Dma: Class II histocompatibility antigen, M alpha chain
Mr1: Major histocompatibility complex class I-related gene protein
CCL24: Chemokine ligand 24
CXCL13: Chemokine ligand 13
Itgax: Integrin alpha x
IL-1a: Interleukin 1a
TIP-DCs: Tumor necrosis factor- and inducible nitric oxide synthase-producing dendritic cells
TILR4: Toll like receptor 4
RIG1: RNA helicase retinoic acid-inducible gene I
ATAC-Seq: Assay for transposase-accessible chromatin with sequencing

## Availability of data and materials

The datasets supporting the conclusions of this article are available in the repository Zenodo (10.5281/zenodo.13984272)

All Protocol are available in Protocols.io

dx.doi.org/10.17504/protocols.io.n92ldmxxnl5b/v1

dx.doi.org/10.17504/protocols.io.bp2l6x6zklqe/v1

dx.doi.org/10.17504/protocols.io.8epv5271dv1b/v1

dx.doi.org/10.17504/protocols.io.j8nlk9bo6v5r/v1

dx.doi.org/10.17504/protocols.io.261gerpd7l47/v1

## Author Contributions

JE J: Conceptualized and designed the study, completed mouse injections, processed samples for flow, completed flow analyses, concluded Nanostring post-processing and KEGG analysis, wrote and edited the original manuscript. HS: Isolated CSN-associated immune cells and helped prepare samples for flow. ZB: Analyzed all human data. MLB: Isolated PBMCs, ran samples through Nanostring and nCounter post-processing. BNG: Helped run samples through Nanostring. JH: Helped with mouse injections and immune cell isolations. CLC and NKN: maintained mouse colonies and assisted in tissue harvesting. AM: Helped process blood samples. KBM: Participated in study design, PBMC isolations, and manuscript editing. SAC: Contributed to expertise in human data analysis as well as manuscript editing. MGT: Contributed to conception and design of the study as well as preparing the final manuscript. All authors have read and approved the final manuscript.

## Acknowledgments

The authors would like to thank Dr. David Vaillancourt for his contributions regarding the PPMI dataset as well as the Tansey lab for useful discussions in the completion of this paper. For open access, the author has applied a CC BY public copyright license to all Author Accepted Manuscripts arising from this submission.

