## Supplemental Figures for "RGS10 Attenuates Systemic Immune Dysregulation Induced by Chronic Inflammatory Stress"

### Slide 1
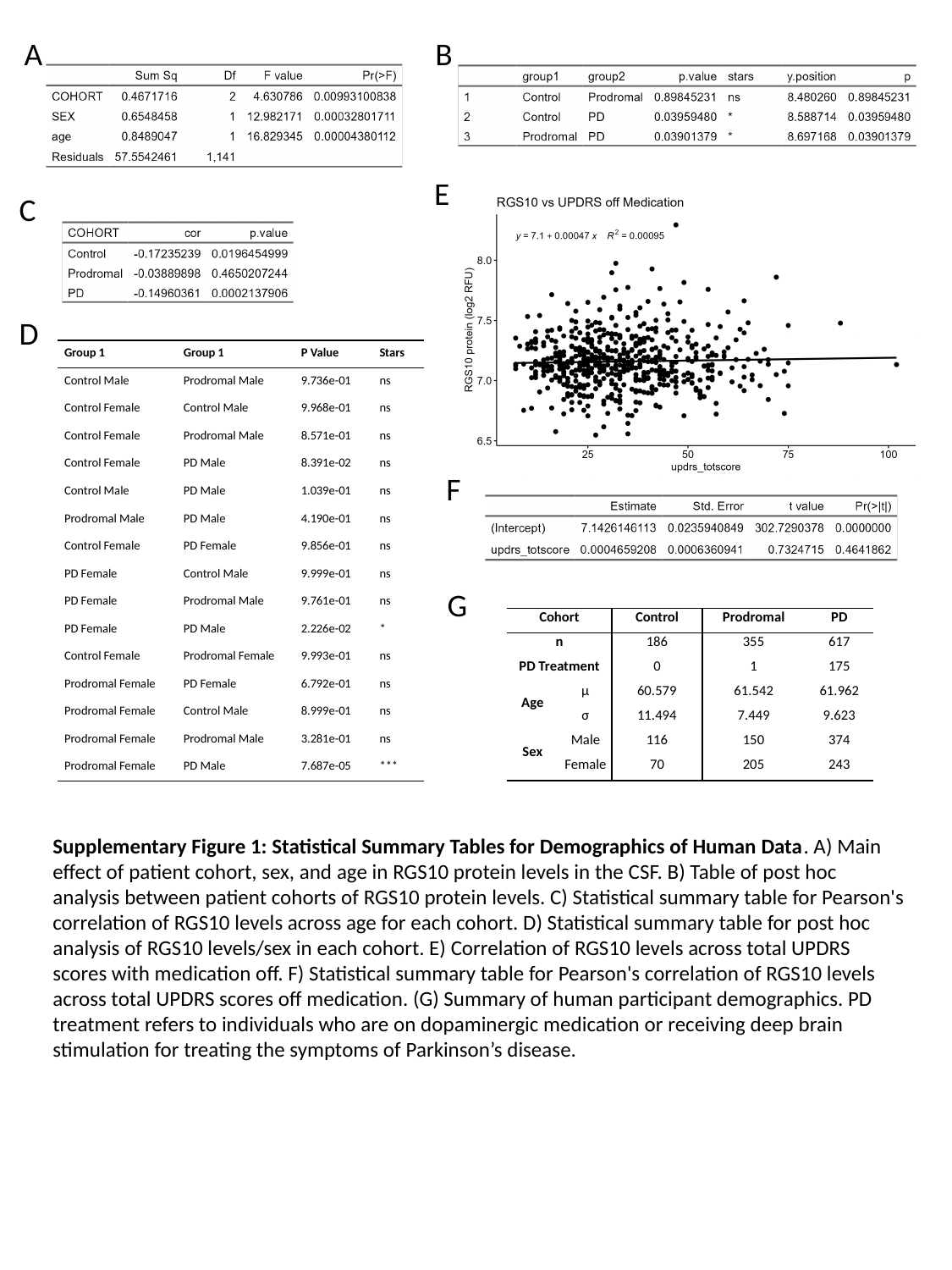

A
B
E
C
D
| Group 1 | Group 1 | P Value | Stars |
| --- | --- | --- | --- |
| Control Male | Prodromal Male | 9.736e-01 | ns |
| Control Female | Control Male | 9.968e-01 | ns |
| Control Female | Prodromal Male | 8.571e-01 | ns |
| Control Female | PD Male | 8.391e-02 | ns |
| Control Male | PD Male | 1.039e-01 | ns |
| Prodromal Male | PD Male | 4.190e-01 | ns |
| Control Female | PD Female | 9.856e-01 | ns |
| PD Female | Control Male | 9.999e-01 | ns |
| PD Female | Prodromal Male | 9.761e-01 | ns |
| PD Female | PD Male | 2.226e-02 | \* |
| Control Female | Prodromal Female | 9.993e-01 | ns |
| Prodromal Female | PD Female | 6.792e-01 | ns |
| Prodromal Female | Control Male | 8.999e-01 | ns |
| Prodromal Female | Prodromal Male | 3.281e-01 | ns |
| Prodromal Female | PD Male | 7.687e-05 | \*\*\* |
F
G
| Cohort | | Control | Prodromal | PD |
| --- | --- | --- | --- | --- |
| n | | 186 | 355 | 617 |
| PD Treatment | | 0 | 1 | 175 |
| Age | μ | 60.579 | 61.542 | 61.962 |
| | σ | 11.494 | 7.449 | 9.623 |
| Sex | Male | 116 | 150 | 374 |
| | Female | 70 | 205 | 243 |
Supplementary Figure 1: Statistical Summary Tables for Demographics of Human Data. A) Main effect of patient cohort, sex, and age in RGS10 protein levels in the CSF. B) Table of post hoc analysis between patient cohorts of RGS10 protein levels. C) Statistical summary table for Pearson's correlation of RGS10 levels across age for each cohort. D) Statistical summary table for post hoc analysis of RGS10 levels/sex in each cohort. E) Correlation of RGS10 levels across total UPDRS scores with medication off. F) Statistical summary table for Pearson's correlation of RGS10 levels across total UPDRS scores off medication. (G) Summary of human participant demographics. PD treatment refers to individuals who are on dopaminergic medication or receiving deep brain stimulation for treating the symptoms of Parkinson’s disease.

### Slide 2
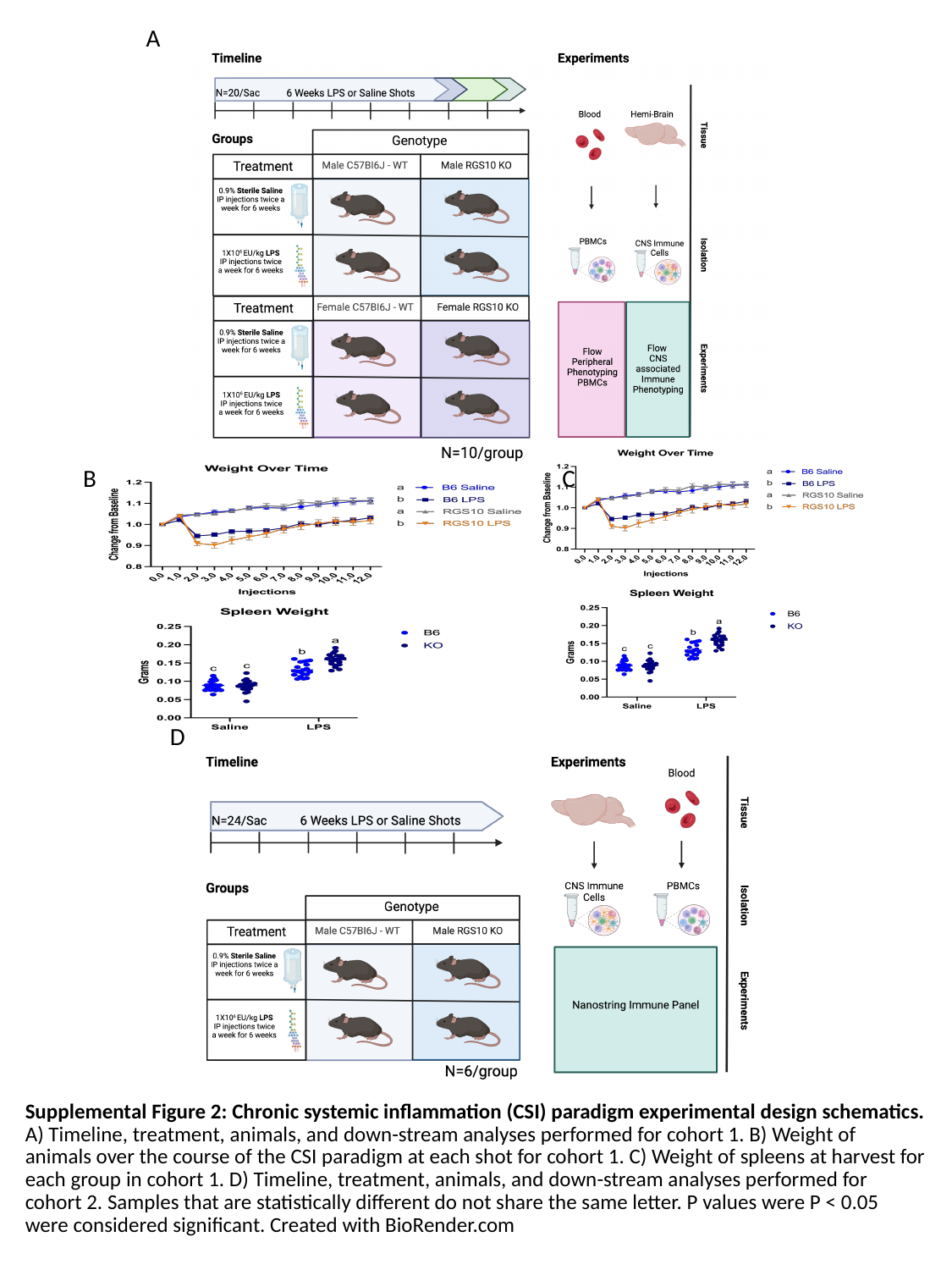

A
C
B
D
Supplemental Figure 2: Chronic systemic inflammation (CSI) paradigm experimental design schematics. A) Timeline, treatment, animals, and down-stream analyses performed for cohort 1. B) Weight of animals over the course of the CSI paradigm at each shot for cohort 1. C) Weight of spleens at harvest for each group in cohort 1. D) Timeline, treatment, animals, and down-stream analyses performed for cohort 2. Samples that are statistically different do not share the same letter. P values were P < 0.05 were considered significant. Created with BioRender.com

### Slide 3
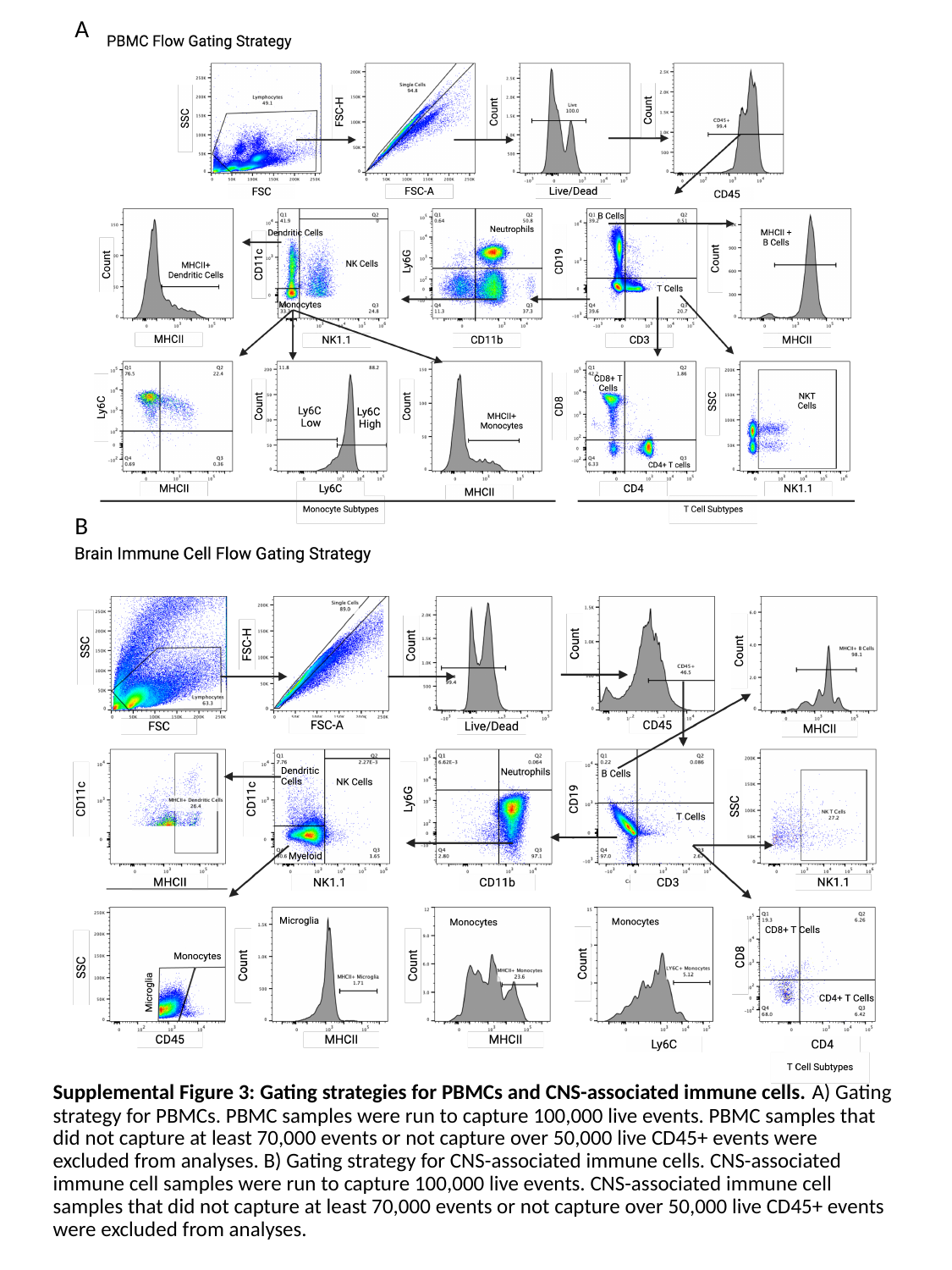

A
B
Supplemental Figure 3: Gating strategies for PBMCs and CNS-associated immune cells. A) Gating strategy for PBMCs. PBMC samples were run to capture 100,000 live events. PBMC samples that did not capture at least 70,000 events or not capture over 50,000 live CD45+ events were excluded from analyses. B) Gating strategy for CNS-associated immune cells. CNS-associated immune cell samples were run to capture 100,000 live events. CNS-associated immune cell samples that did not capture at least 70,000 events or not capture over 50,000 live CD45+ events were excluded from analyses.

### Slide 4
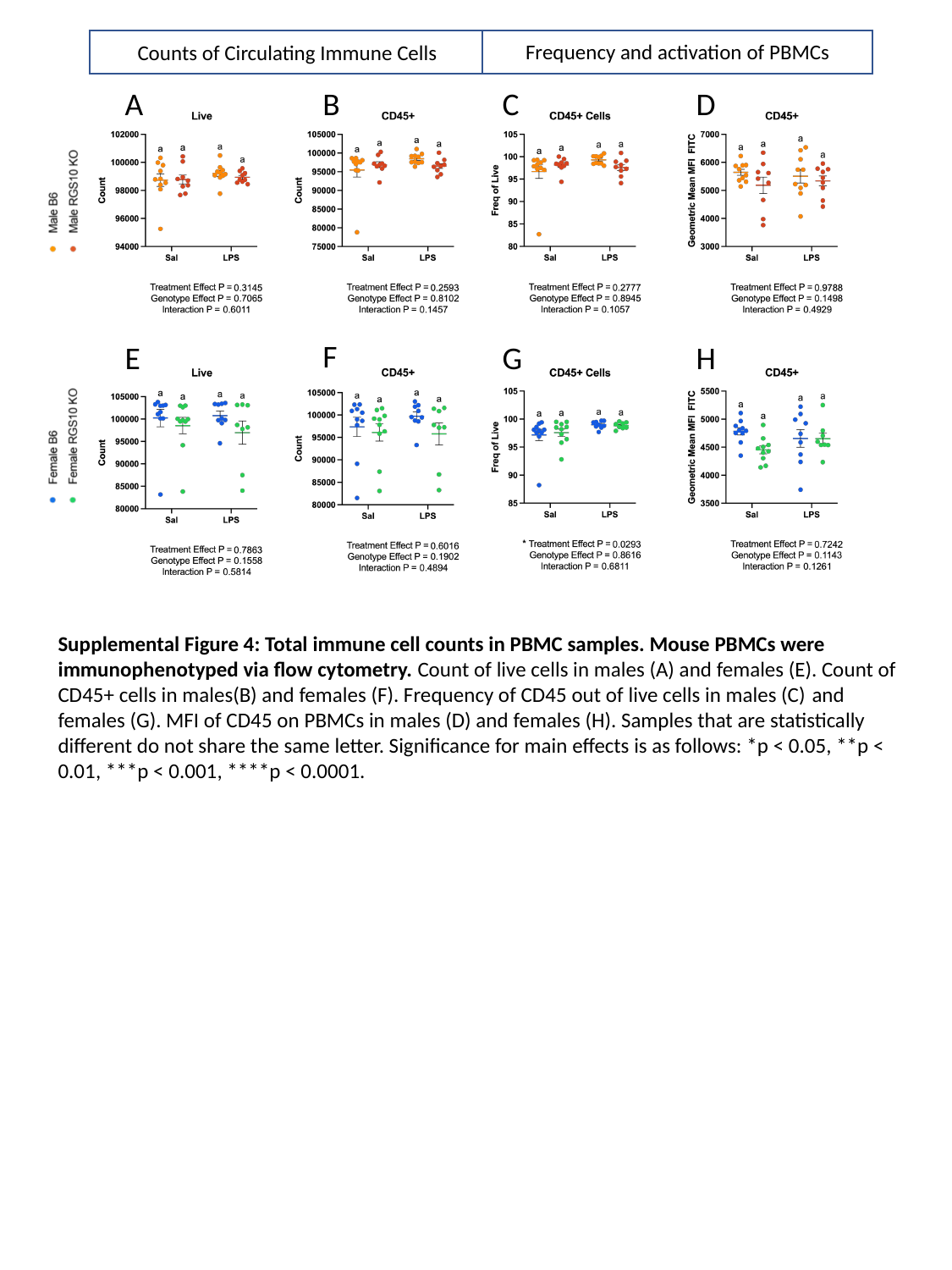

Frequency and activation of PBMCs
Counts of Circulating Immune Cells
B
A
C
D
F
E
G
H
Supplemental Figure 4: Total immune cell counts in PBMC samples. Mouse PBMCs were immunophenotyped via flow cytometry. Count of live cells in males (A) and females (E). Count of CD45+ cells in males(B) and females (F). Frequency of CD45 out of live cells in males (C) and females (G). MFI of CD45 on PBMCs in males (D) and females (H). Samples that are statistically different do not share the same letter. Significance for main effects is as follows: *p < 0.05, **p < 0.01, ***p < 0.001, ****p < 0.0001.

### Slide 5
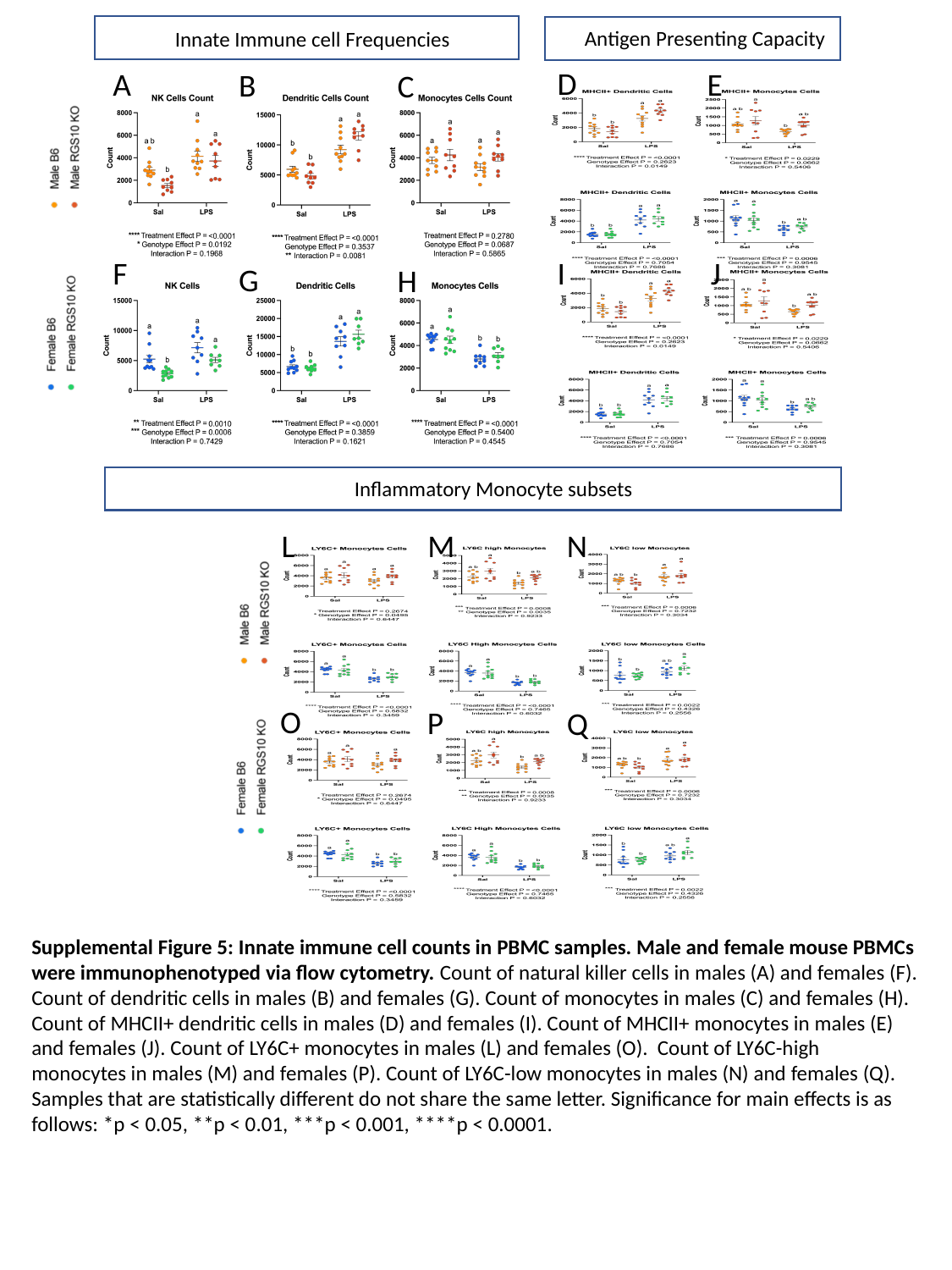

Antigen Presenting Capacity
D
E
I
J
Innate Immune cell Frequencies
A
B
C
F
G
H
Inflammatory Monocyte subsets
M
N
L
O
P
Q
Supplemental Figure 5: Innate immune cell counts in PBMC samples. Male and female mouse PBMCs were immunophenotyped via flow cytometry. Count of natural killer cells in males (A) and females (F). Count of dendritic cells in males (B) and females (G). Count of monocytes in males (C) and females (H). Count of MHCII+ dendritic cells in males (D) and females (I). Count of MHCII+ monocytes in males (E) and females (J). Count of LY6C+ monocytes in males (L) and females (O). Count of LY6C-high monocytes in males (M) and females (P). Count of LY6C-low monocytes in males (N) and females (Q). Samples that are statistically different do not share the same letter. Significance for main effects is as follows: *p < 0.05, **p < 0.01, ***p < 0.001, ****p < 0.0001.

### Slide 6
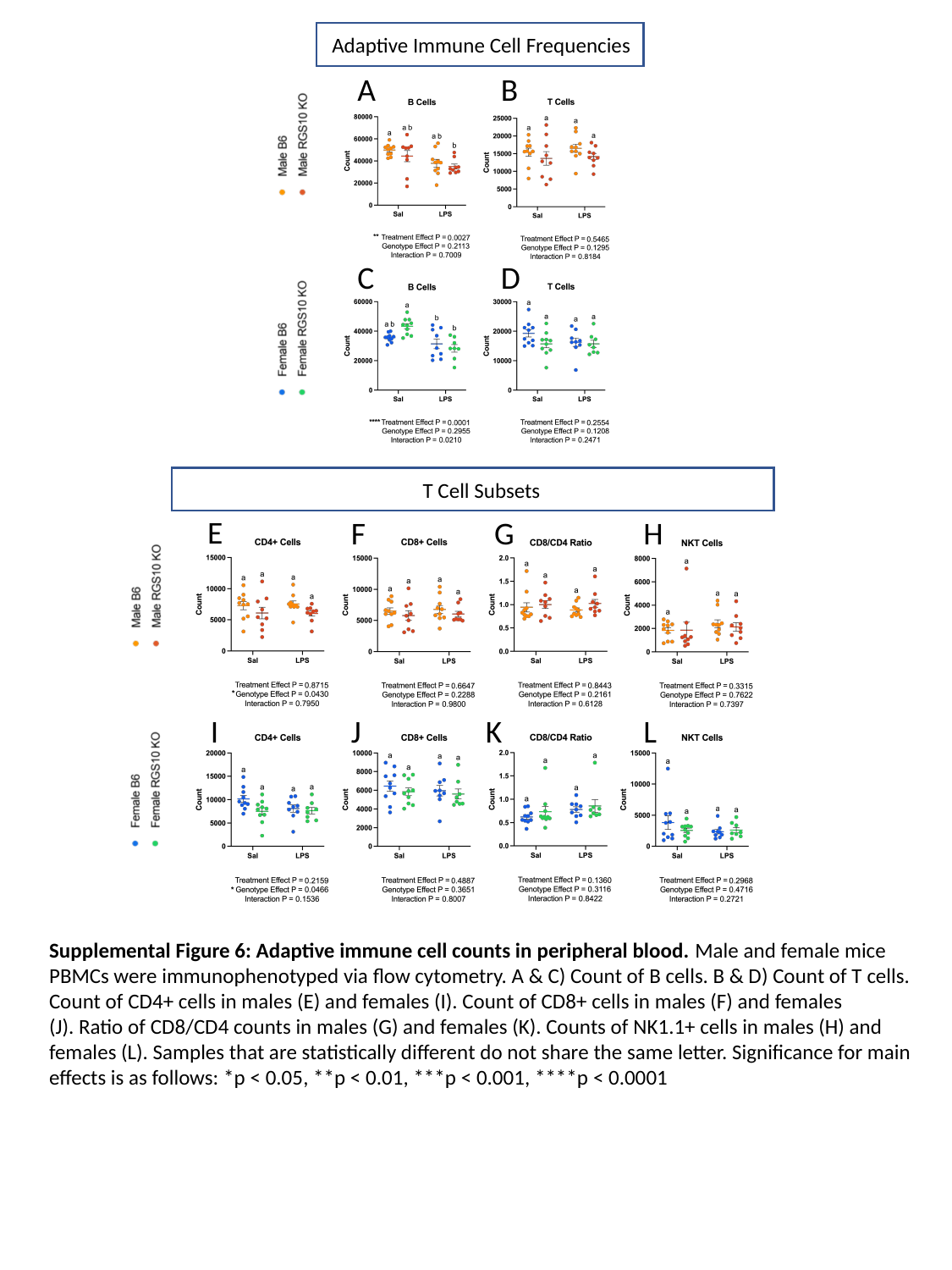

Adaptive Immune Cell Frequencies
A
B
C
D
T Cell Subsets
E
F
G
H
I
J
K
L
Supplemental Figure 6: Adaptive immune cell counts in peripheral blood. Male and female mice PBMCs were immunophenotyped via flow cytometry. A & C) Count of B cells. B & D) Count of T cells. Count of CD4+ cells in males (E) and females (I). Count of CD8+ cells in males (F) and females (J). Ratio of CD8/CD4 counts in males (G) and females (K). Counts of NK1.1+ cells in males (H) and females (L). Samples that are statistically different do not share the same letter. Significance for main effects is as follows: *p < 0.05, **p < 0.01, ***p < 0.001, ****p < 0.0001

### Slide 7
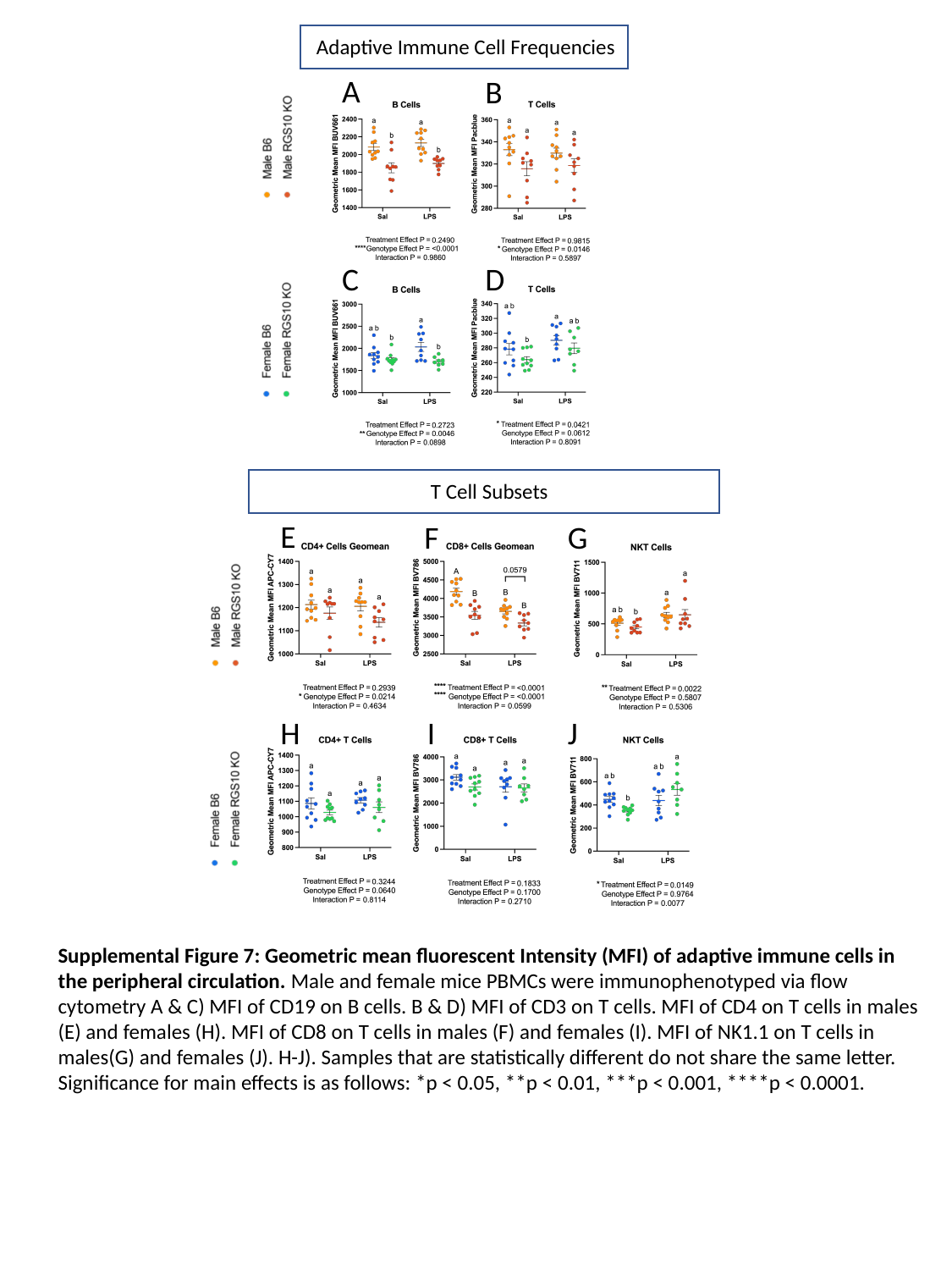

Adaptive Immune Cell Frequencies
A
B
C
D
T Cell Subsets
E
F
G
H
I
J
Supplemental Figure 7: Geometric mean fluorescent Intensity (MFI) of adaptive immune cells in the peripheral circulation. Male and female mice PBMCs were immunophenotyped via flow cytometry A & C) MFI of CD19 on B cells. B & D) MFI of CD3 on T cells. MFI of CD4 on T cells in males (E) and females (H). MFI of CD8 on T cells in males (F) and females (I). MFI of NK1.1 on T cells in males(G) and females (J). H-J). Samples that are statistically different do not share the same letter. Significance for main effects is as follows: *p < 0.05, **p < 0.01, ***p < 0.001, ****p < 0.0001.

### Slide 8
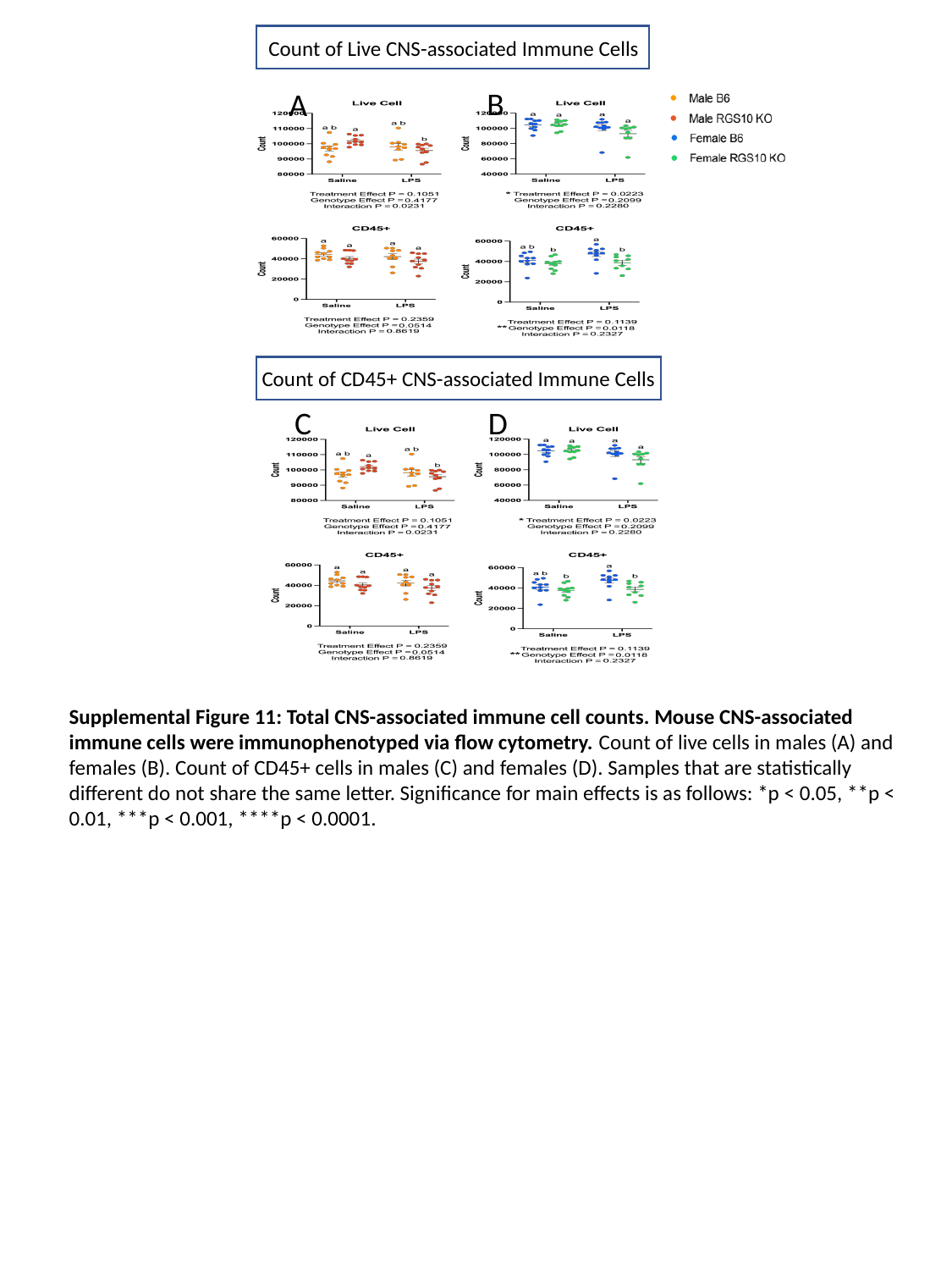

Count of Live CNS-associated Immune Cells
B
A
Count of CD45+ CNS-associated Immune Cells
C
D
Supplemental Figure 11: Total CNS-associated immune cell counts. Mouse CNS-associated immune cells were immunophenotyped via flow cytometry. Count of live cells in males (A) and females (B). Count of CD45+ cells in males (C) and females (D). Samples that are statistically different do not share the same letter. Significance for main effects is as follows: *p < 0.05, **p < 0.01, ***p < 0.001, ****p < 0.0001.

### Slide 9
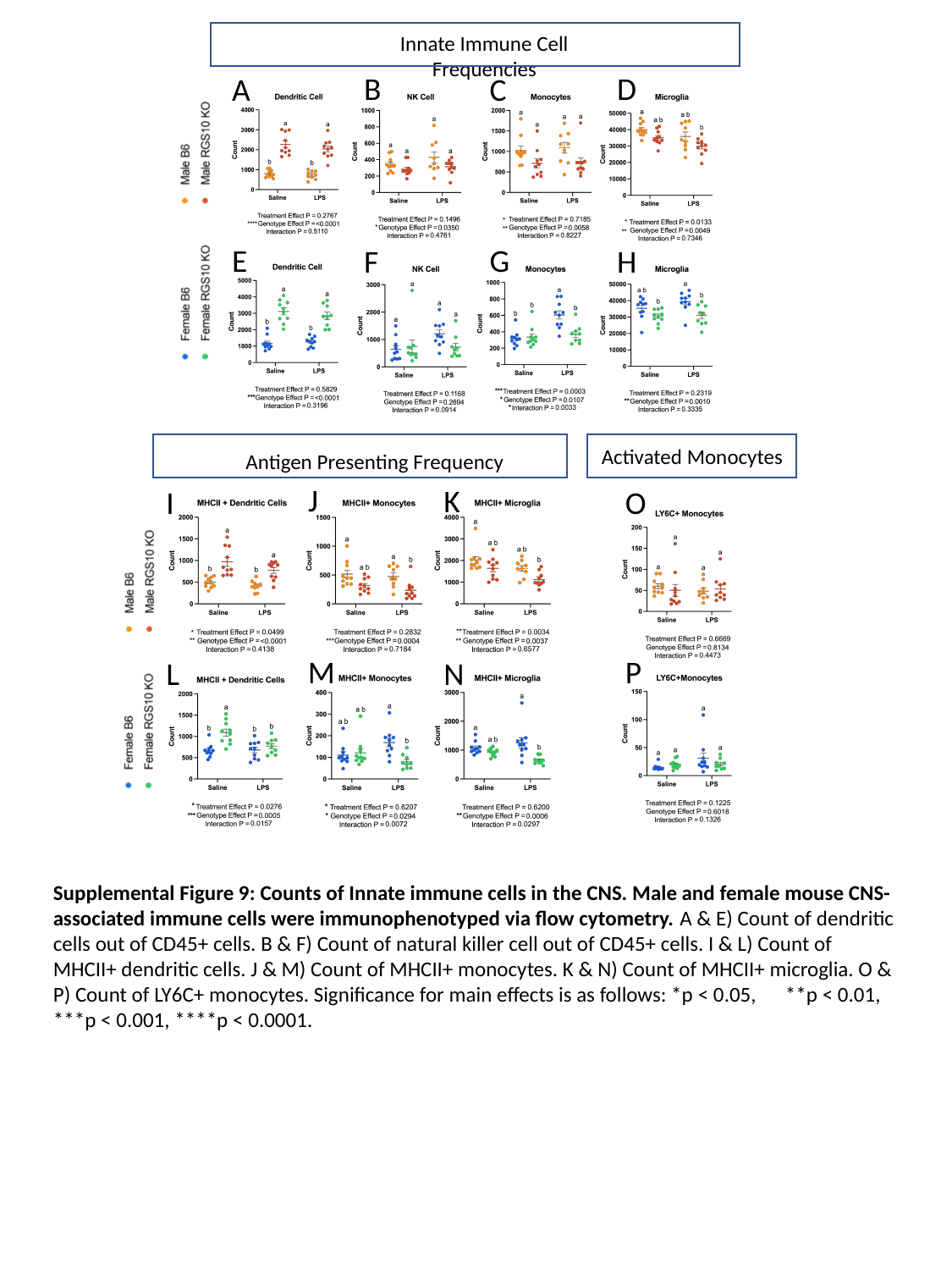

Innate Immune Cell Frequencies
D
B
A
C
G
E
H
F
Activated Monocytes
Antigen Presenting Frequency
J
K
I
O
M
P
L
N
Supplemental Figure 9: Counts of Innate immune cells in the CNS. Male and female mouse CNS-associated immune cells were immunophenotyped via flow cytometry. A & E) Count of dendritic cells out of CD45+ cells. B & F) Count of natural killer cell out of CD45+ cells. I & L) Count of MHCII+ dendritic cells. J & M) Count of MHCII+ monocytes. K & N) Count of MHCII+ microglia. O & P) Count of LY6C+ monocytes. Significance for main effects is as follows: *p < 0.05, **p < 0.01, ***p < 0.001, ****p < 0.0001.

### Slide 10
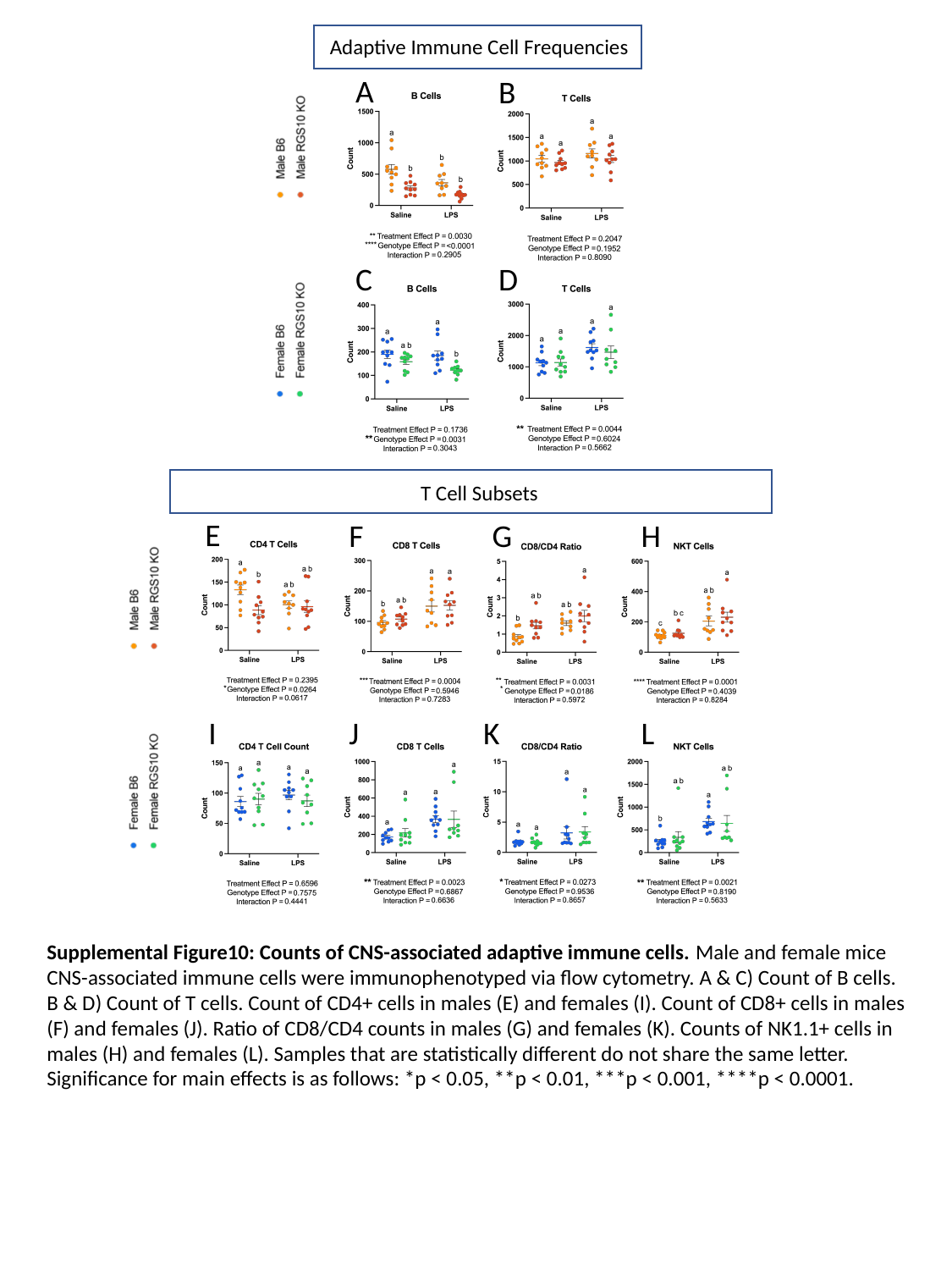

Adaptive Immune Cell Frequencies
A
B
C
D
T Cell Subsets
E
F
G
H
I
J
K
L
Supplemental Figure10: Counts of CNS-associated adaptive immune cells. Male and female mice CNS-associated immune cells were immunophenotyped via flow cytometry. A & C) Count of B cells. B & D) Count of T cells. Count of CD4+ cells in males (E) and females (I). Count of CD8+ cells in males (F) and females (J). Ratio of CD8/CD4 counts in males (G) and females (K). Counts of NK1.1+ cells in males (H) and females (L). Samples that are statistically different do not share the same letter. Significance for main effects is as follows: *p < 0.05, **p < 0.01, ***p < 0.001, ****p < 0.0001.

### Slide 11
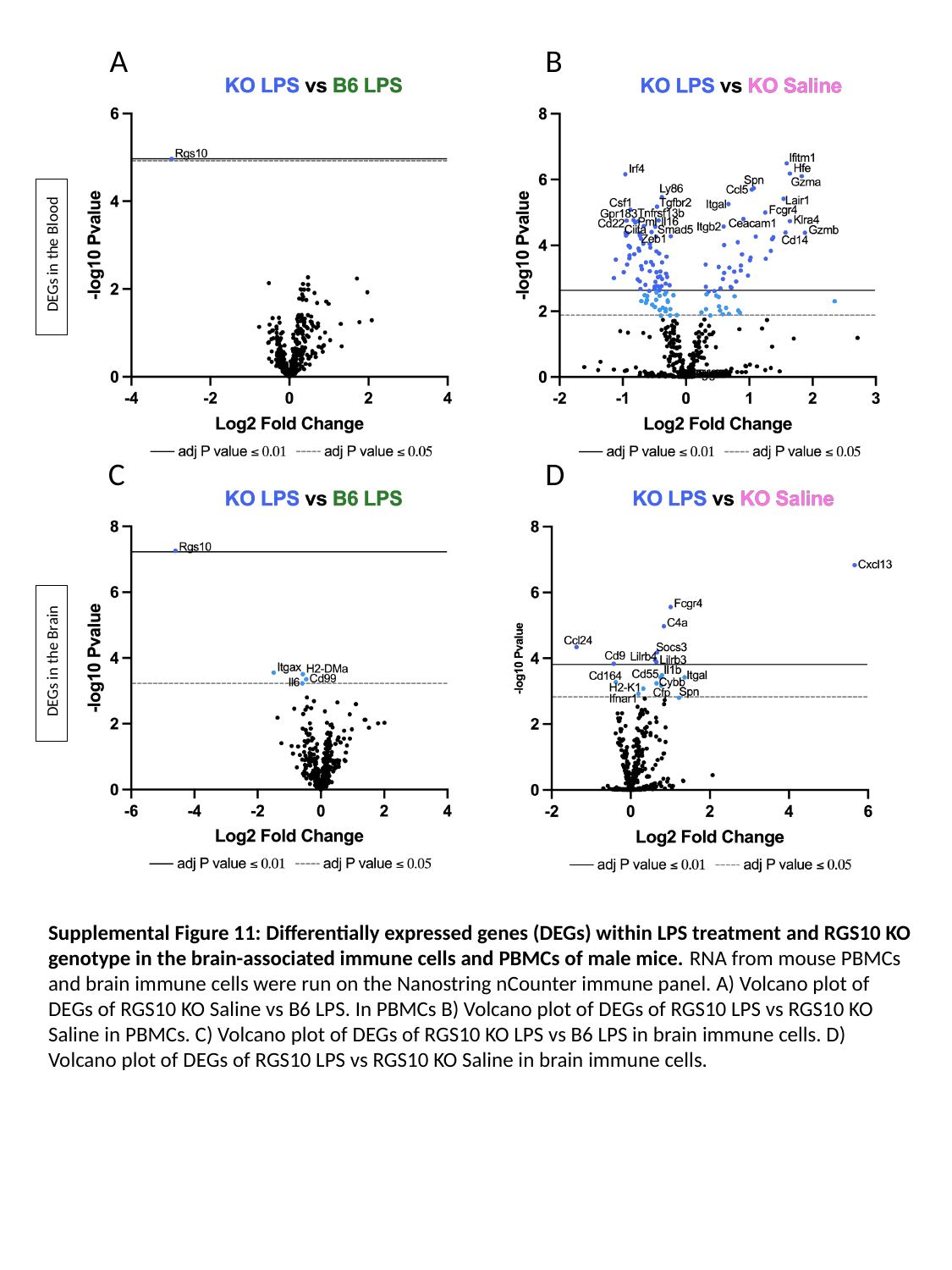

A
B
 DEGs in the Blood
C
D
 DEGs in the Brain
Supplemental Figure 11: Differentially expressed genes (DEGs) within LPS treatment and RGS10 KO genotype in the brain-associated immune cells and PBMCs of male mice. RNA from mouse PBMCs and brain immune cells were run on the Nanostring nCounter immune panel. A) Volcano plot of DEGs of RGS10 KO Saline vs B6 LPS. In PBMCs B) Volcano plot of DEGs of RGS10 LPS vs RGS10 KO Saline in PBMCs. C) Volcano plot of DEGs of RGS10 KO LPS vs B6 LPS in brain immune cells. D) Volcano plot of DEGs of RGS10 LPS vs RGS10 KO Saline in brain immune cells.

### Slide 12
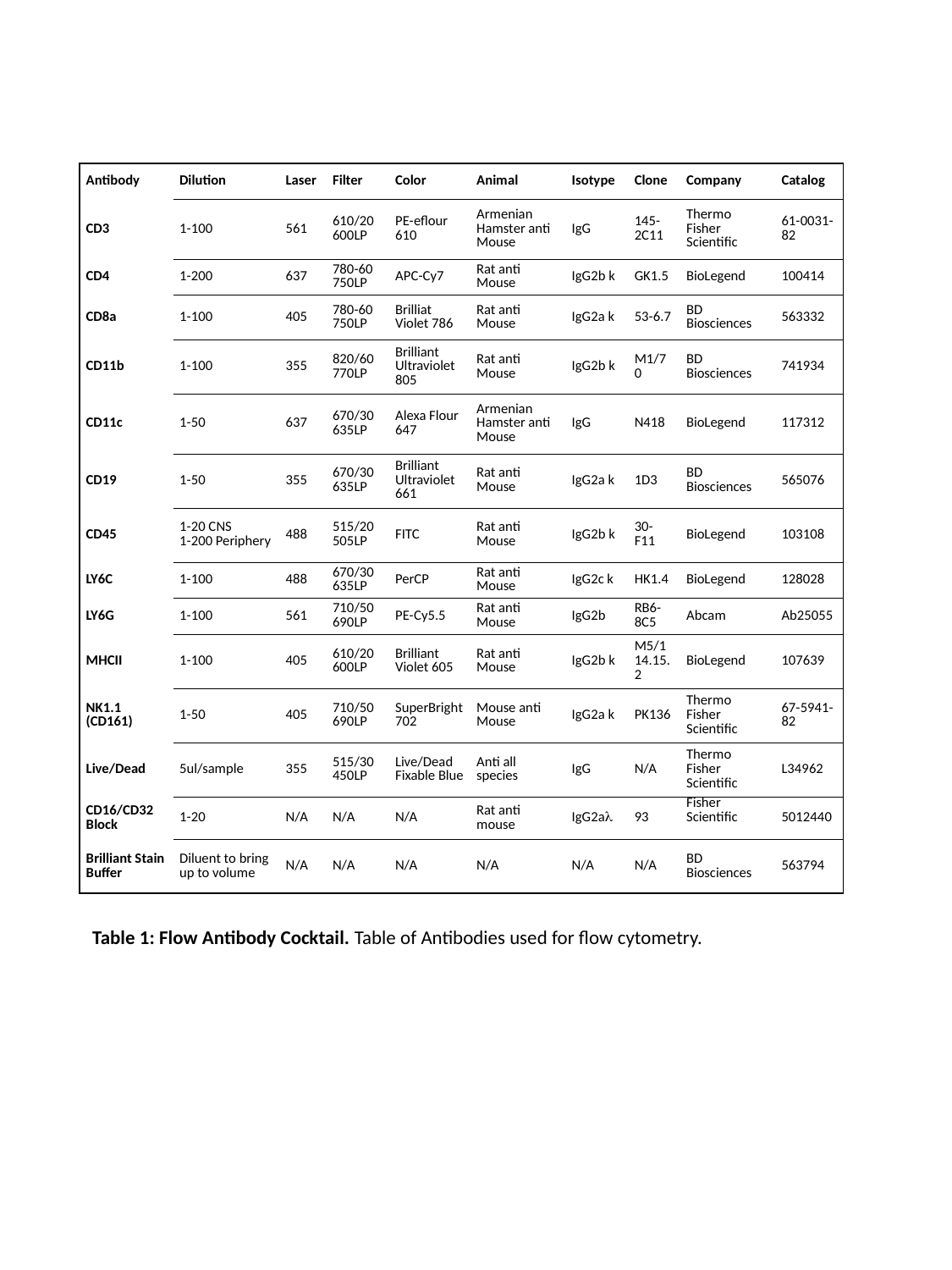

| Antibody | Dilution | Laser | Filter | Color | Animal | Isotype | Clone | Company | Catalog |
| --- | --- | --- | --- | --- | --- | --- | --- | --- | --- |
| CD3 | 1-100 | 561 | 610/20 600LP | PE-eflour 610 | Armenian Hamster anti Mouse | IgG | 145-2C11 | Thermo Fisher Scientific | 61-0031-82 |
| CD4 | 1-200 | 637 | 780-60 750LP | APC-Cy7 | Rat anti Mouse | IgG2b k | GK1.5 | BioLegend | 100414 |
| CD8a | 1-100 | 405 | 780-60 750LP | Brilliat Violet 786 | Rat anti Mouse | IgG2a k | 53-6.7 | BD Biosciences | 563332 |
| CD11b | 1-100 | 355 | 820/60 770LP | Brilliant Ultraviolet 805 | Rat anti Mouse | IgG2b k | M1/70 | BD Biosciences | 741934 |
| CD11c | 1-50 | 637 | 670/30 635LP | Alexa Flour 647 | Armenian Hamster anti Mouse | IgG | N418 | BioLegend | 117312 |
| CD19 | 1-50 | 355 | 670/30 635LP | Brilliant Ultraviolet 661 | Rat anti Mouse | IgG2a k | 1D3 | BD Biosciences | 565076 |
| CD45 | 1-20 CNS 1-200 Periphery | 488 | 515/20 505LP | FITC | Rat anti Mouse | IgG2b k | 30-F11 | BioLegend | 103108 |
| LY6C | 1-100 | 488 | 670/30 635LP | PerCP | Rat anti Mouse | IgG2c k | HK1.4 | BioLegend | 128028 |
| LY6G | 1-100 | 561 | 710/50 690LP | PE-Cy5.5 | Rat anti Mouse | IgG2b | RB6-8C5 | Abcam | Ab25055 |
| MHCII | 1-100 | 405 | 610/20 600LP | Brilliant Violet 605 | Rat anti Mouse | IgG2b k | M5/114.15.2 | BioLegend | 107639 |
| NK1.1 (CD161) | 1-50 | 405 | 710/50 690LP | SuperBright 702 | Mouse anti Mouse | IgG2a k | PK136 | Thermo Fisher Scientific | 67-5941-82 |
| Live/Dead | 5ul/sample | 355 | 515/30 450LP | Live/Dead Fixable Blue | Anti all species | IgG | N/A | Thermo Fisher Scientific | L34962 |
| CD16/CD32 Block | 1-20 | N/A | N/A | N/A | Rat anti mouse | IgG2aλ | 93 | Fisher Scientific | 5012440 |
| Brilliant Stain Buffer | Diluent to bring up to volume | N/A | N/A | N/A | N/A | N/A | N/A | BD Biosciences | 563794 |
Table 1: Flow Antibody Cocktail. Table of Antibodies used for flow cytometry.
